# Metal ion activation and DNA recognition by the *Deinococcus radiodurans* manganese sensor DR2539

**DOI:** 10.1101/2024.02.12.579695

**Authors:** Cristiano Mota, Myles Webster, Melissa Saidi, Ulrike Kapp, Chloe Zubieta, Gabriele Giachin, José Antonio Manso, Daniele de Sanctis

## Abstract

The accumulation of manganese ions is crucial for scavenging reactive oxygen species (ROS) and protecting the proteome of *Deinococcus radiodurans* (*Dr*). However, metal homeostasis still needs to be tightly regulated to avoid toxicity. DR2539, a dimeric transcription regulator, plays a key role in *Dr* manganese homeostasis. Despite comprising three well-conserved domains: a DNA binding domain, a dimerization domain, and an ancillary domain, both the metal ion activation mechanism and the DNA recognition mechanism remain elusive. In this study, we present biophysical analyses and the structure of the dimerization and DNA binding domains of DR2539 in its holo form and in complex with the 21 bp pseudo-palindromic repeat of the *dr1709* promotor region. These findings shed light into the activation and recognition mechanisms. The dimer presents eight manganese binding sites that induce structural conformations essential for DNA binding. The analysis of the protein-DNA interfaces elucidates the significance of Tyr59 and helix H3 sequence in the interaction with the DNA. Finally, the structure in solution as determined by small angle X-ray scattering experiments and supported by AlphaFold modelling provides a model illustrating the conformational changes induced upon metal binding.

## Introduction

*Deinococcus radiodurans* (*Dr*) is a gram-positive bacterium highly resistant to stress conditions [1]. Since its discovery in 1956 [2], the molecular basis of its resistance has been the subject of several studies. While distinctive features such as multiple copies of its genome [3] and an efficient DNA repair system capable of restoring the entire genome through homologous recombination have been identified [4–7], these characteristics alone do not explain the resistance of an organism to environmental challenges. Beyond this, *Dr* presents a unique ability to protect its proteome against oxidative damage induced by reactive oxygen species (ROS). This exceptional trait enables *Dr* proteins to remain fully functional after exposure to stress conditions, and facilitates the recovery and repair of bacterial DNA [8,9].

*Dr* presents higher ROS scavenging capacity when compared with other bacteria [10,11]. The scavenging is mediated by enzymatic (e.g. catalases, superoxide dismutases, and peroxidases) and non-enzymatic (e.g. divalent manganese complexes and carotenoids) components. Among these, manganese ions complexed with phosphates, nucleotides and amino acids emerged as the most powerful ROS scavengers in this bacterium [11]. The cellular concentration of these ions in *Dr* is higher compared with other bacteria, in contrast with relatively low iron levels[8]. However if the high content of manganese benefits ROS scavenging, an excess of this divalent ion is still toxic to the bacteria and the manganese/iron homeostasis needs to be strictly regulated. Within the *Dr* genome, three type of manganese-dependent transport genes have been annotated: *dr1236* (encoding a manganese efflux protein) [12], *dr1709* (encoding an Nramp family transporter) [13–15], and the genes encoding ATP-dependent transporters (*dr2283*, *dr2284* and *dr2523*) [14,16,17]. On the other hand, the genes that are involved in iron-dependent transport encode an ABC-type hemin transporter (*drb0016*), an ABC-type Fe(III)-siderophore transporter (*drb0017*), two Fe(II) transporters (*dr1219*, *dr1120*) and two DNA protection proteins (Dps) (*dr2263*, *drb0092*) [18]. The regulation of these transport systems is managed by three oxidation-related regulators, OxyR (DR0615) [19], the Fur homolog, DR0865 [20] and DR2539 [16,17]. The DR0615 protein is both a transcriptional activator of the *katE* and *drb0125* genes and a transcriptional repressor of the *dps* and *mntH* genes. The Fur homologue, DR0865, is a positive regulator of *dr1236* efflux manganese-transporter and a negative regulator of *dr2283*, *dr2284* and *dr2523* transporters [20]. DR2539 is reported to be a down-regulator of manganese transporter genes (*dr1709* and *dr2283*) and an up-regulator of iron-dependent transporter genes (*dr1219* and *drb0125*) [17]. DR1709 is a member of the divalent metal transporters Nramp (natural-associated macrophage protein) family [15]. Transporters that belong to the Nramp family have been identified in organisms ranging from bacteria to humans, with a preference for transporting iron among eukaryotes and manganese in prokaryotic species [21–26]. This manganese transporter has been revealed to be essential for *Dr* growth [14] and one of the key proteins governing the remarkable manganese/iron homeostasis in the bacterium. Due to its regulatory role in *dr1709*, DR2539 has been proposed as the principal regulator of intracellular manganese homeostasis in *Dr* [16]. The diphtheria toxin regulator (DtxR) family, to which DR2539 belongs, was initially described with *Corynebacterium diphtheriae* DtxR (*Cd*DtxR) as an iron-responsive repressor [27]. Later, extensive sequencing of bacterial genomes has revealed the details of multiple DtxR-like repressors, including IdeR from *Mycobacterium tuberculosis* (*Mt*IdeR) [28,29], *Thermoplasma acidophilum* (*Ta*IdeR) [30] and *Sachararopolyspora erythraea* (*Se*IdeR) [31], ScaR from *Streptococus gordonii* (*Sg*Scar) [32], MntR from *Staphilococcus aureus* (*Sa*MntR) *Bacillus subtilis* (*Bs*MntR), *Bacillus halodurans* (*Bh*MntR) and *Mycobacterium tuberculosis* (*Mt*MntR) [33–38], TroR from *Treponema pallidum* (*Tp*TroR) [39], MtsR from *Streptococcus pyrogenes* (*Sp*MtsR) [40,41], SirR *Staphylococcus epidermidis* (*Se*SirR) [42], and SloR from *Streptococcus mutans* (*Sm*SloR) [43–45]. Several structures of *Cd*DtxR homologues have been solved, revealing a modular domain organisation including an N-terminal DNA-binding domain (DBD), a dimerization domain (DD) and a C-terminal domain, the ancillary domain (AD). Notably, the AD is missing in some MntRs. The molecular mechanisms of metal- and DNA-binding were already proposed by several authors [46–48], but structural differences and the variation in the number of metal-binding sites suggest different types of regulation among the different homologues [32].

In this study, we performed biochemical and biophysical studies to characterise the DR2539 regulator. We solved the crystal structures of the DNA binding and dimerization domains in the holo-form, and in complex with the *dr1709* promotor region with its physiological cofactor (manganese). In addition to the primary and ancillary metal binding sites, observed in other DtxR regulators, DR2539 reveals a novel metal center. Furthermore, SAXS studies, supported by the predicted model, contributed additional insights into the metal-induced conformational changes, the ancillary domain, and the full-length arrangement. These structures provide insights into the metal sensing and DNA binding mechanisms of this key regulator of manganese/iron homeostasis in *Dr*.

## Results

### DR2539 expression and purification

*dr2539* was cloned directly from *Dr* and inserted into an *E. coli* expression plasmid with an N-terminal 6x histidine tag. The recombinant DR2539 protein was overexpressed and purified using Ni (II) affinity chromatography. During the initial purification step, it was possible to observe two species that corresponded to the full length (25 kDa) and a cleaved stable form with 16 kDa molecular weight. Both species were confirmed by N-terminal sequencing and the mass determined by mass spectrometry (ESI-TOF), indicating that the protein suffered proteolysis in the C-terminus before the first purification step. Both forms were well-separated using heparin affinity column purification (Figure S1).

### Overall structure of truncated DR2539 in the holo-form

The crystal structure of truncated DR2539 (DR2539tr, residues 4-142) was determined by SAD phasing using a dataset collected at 2.5 Å resolution at the wavelength of 1.771 Å. DR2539 crystallised in the P4_3_2_1_2 space group with unit cell dimensions a= b= 60.8 Å, c= 187.7 Å. The anomalous signal from Cd, present in the crystallisation buffer, allowed to locate a total of 17 anomalous peaks that were used to calculate the experimental phases. The refined model (R_work_ and R_free_ values of 20.5% and 25.5%, Table S1, PDB ID: 8PVT) was used for molecular replacement to solve the structure of a “native” dataset, collected at 2.0 Å resolution at the wavelength of 0.980 Å, which was eventually refined to crystallographic R_work_ and R_free_ values of 18.6% and 22.0%, respectively, with good geometry. This refined model (PDB ID 8PVZ) contains 271 amino acid residues from two monomers in the asymmetric unit. The biological dimer is obtained by a symmetric operation and presents approximate dimensions of 65.7 x 46.6 x 35.5 Å (Figure 1). Each subunit is divided into three domains, a N-terminal DNA binding domain (DBD, residues 4-64), a dimerization domain (DD, residues 65 –127) and a coil region (residues 128-141) that links the missing ancillary domain (Figure 1 and S2). The DBD comprises three α-helices and a region that corresponds to a pair of antiparallel β-strands forming a hairpin, constituting a winged helix-turn-helix (HTH) motif. The region corresponding to the wing, antiparallel β-strands, appears to be highly flexible, as evidenced by the higher B-factors. In one of the protomers, this region could not be fully modelled. The DBD and the DD are connected via α4 (residues 65-91) linking the DNA recognition domain to α5 and α6 helices from the DD. The solvent-accessible surface buried at the interface between the DD domains of the two monomers is 840.7 Å^2^ (∼ 11% of the monomer surface area), with 21 amino acid residues involved in this interface (PDBePISA protein-protein interaction server; http://www.ebi.ac.uk/msd-srv/prot_int/). The dimer interface is mainly constituted by a hydrophobic patch composed by Leu82, Phe85, Leu86, Ala89, Leu90, Val92, Pro93, Ala103, Leu104, Ala107, Leu108, Leu112, Ile116 and Trp119 side chains, and by a network of thirteen hydrogen bonds between and eleven salt bridges (Table S2). Gel filtration profiles of DR2539 incubated with manganese ions or in presence of EDTA yielded a single peak without significant differences between the elution profiles (Figure S3) suggesting that DR2539 exists as a functional dimer in solution either in the apo or holo form.

**Figure 1.**
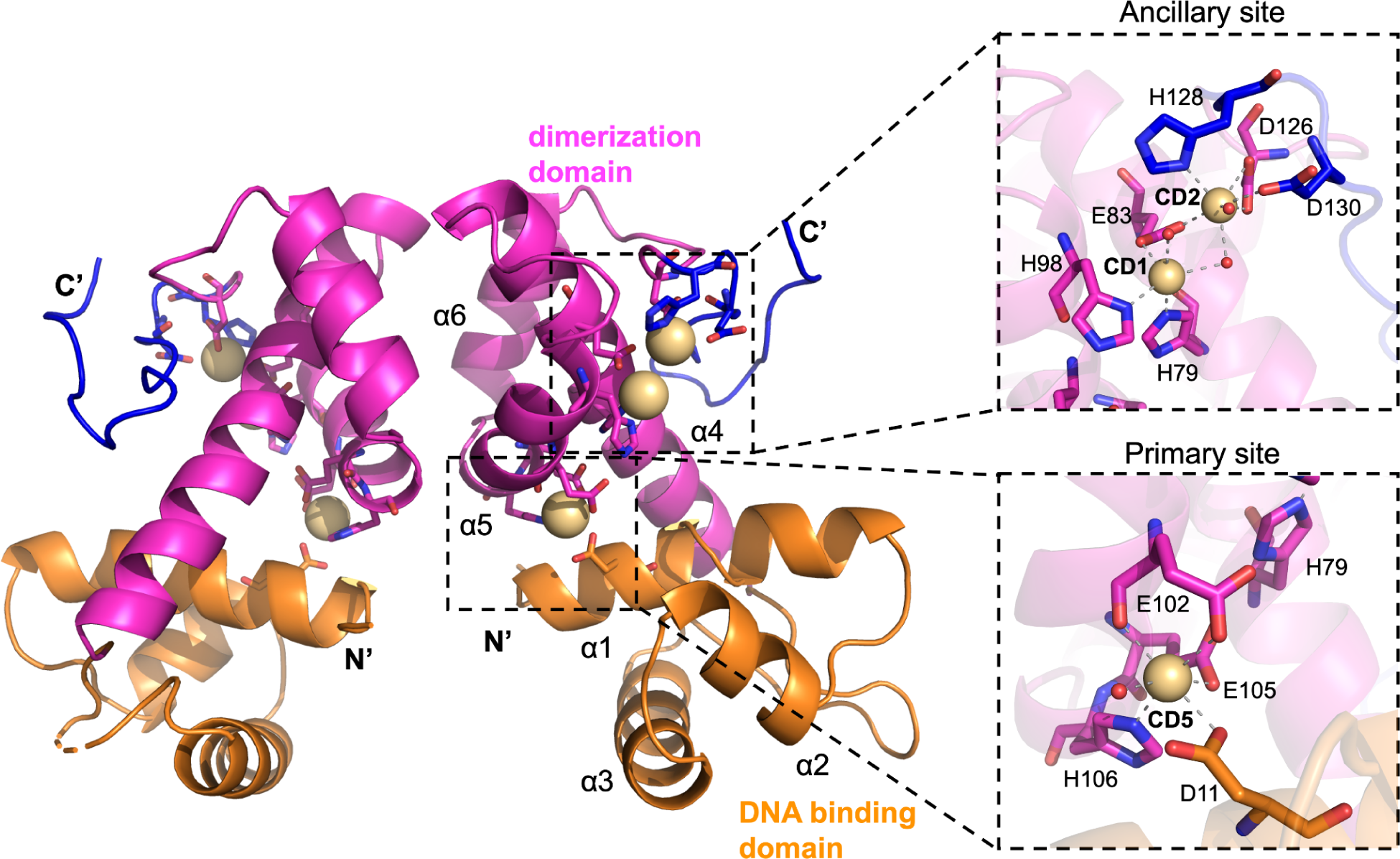
The three-dimensional structure of holo DR2539tr. On the left, the biological dimer, generated by a two fold crystallographic symmetry operation. On the right, close up views of the ancillary (top) and primary (bottom) metal binding sites. In orange the DNA binding domain, in magenta de dimerization domain and in blue the linker that would connect the missing c-terminal ancillary domain. Cadmium atoms are represented as spheres in wheat colour.

Cadmium ions present in the crystallisation buffer were used to calculate the experimental phases and provided initial insights into DR2539 metal binding sites. The cadmiums bound to the protein were confirmed in the anomalous difference map calculated from the high resolution “native” dataset, which presents 12 peaks higher than 10 σ and 4 additional peaks at about 5 σ. Although, 22 Cd^2+^ ions were modelled in the final model, only 4 metal-binding sites were refined with full occupancy, two for each monomer, with temperature B-factors of 34.1 Å^2^, 34.6 Å^2^, 33.1 Å^2^ and 35.2 Å^2^, for CD1, CD2, CD3 and CD4, respectively. These four ions are bound to the two ancillary sites present in the asymmetric unit (Figure 1) and they form binuclear metal centres. Two other cadmium ions are bound to the two primary sites present in the asymmetric unit (CD5 and CD8) with occupancies refined to 0.82 and 0.83 and B-factor of 45.6 Å^2^ and 33.3 Å^2^, respectively. Additionally, a previously unreported binding site is observed between α4 and α6 with high occupancies of 0.89 and 0.92 and B-factor of 36.1 Å^2^ and 34.9 Å^2^, respectively for the two chains (CD7 and CD9) (Table S3). This putative new metal binding site is hexacoordinated to Glu74OE1, His78NE2, Glu113OE1, a bidentate interaction to Glu110 and one water molecule. The remaining cadmium ions are solvent exposed and are coordinated by charged residues, His15NE2, Glu50OE2, His56NE2, His87NE2, Asp95OD2, Asp99OD1 and His125ND1with partial occupancies (Table S3) that are lower than the other sites and are most likely due to the high concentration of cadmium in the crystallisation solution.

### Overall structure of DR2539tr-*dr1709p*-manganese complex

DR2539 specifically recognizes the promotor region (ATTTTAGTCGCGCCTAAAAT) of its target gene (*dr1709*) and affects its transcription [16,17]. We obtained crystals of DR2539tr bound to the 21bp promoter region of the manganese transporter *dr1709* gene (*dr1709p*) and we solved the structure of the complex at 2.2 Å by molecular replacement using the structure of the holo form (PDB ID 8PVZ) as the search model. This structure, refined to crystallographic R_work_ and R_free_ values of 18.3% and 22.7%, (Table S1) contains one protein monomer of 136 amino acids and a 21-mer single strand DNA. Because the DNA sequence is quasi-palindromic (differing in 2 base pairs) the 21-mer double stranded DNA was refined with 50% occupancy in the asymmetric unit with the biological dimer generated by a crystallographic two-fold axis (Figure 2A). For this reason and because of the odd number of nucleotides there is ambiguity in three base pairs (forward Thy8, Cyt13, and Ade21, Figure S4).

**Figure 2.**
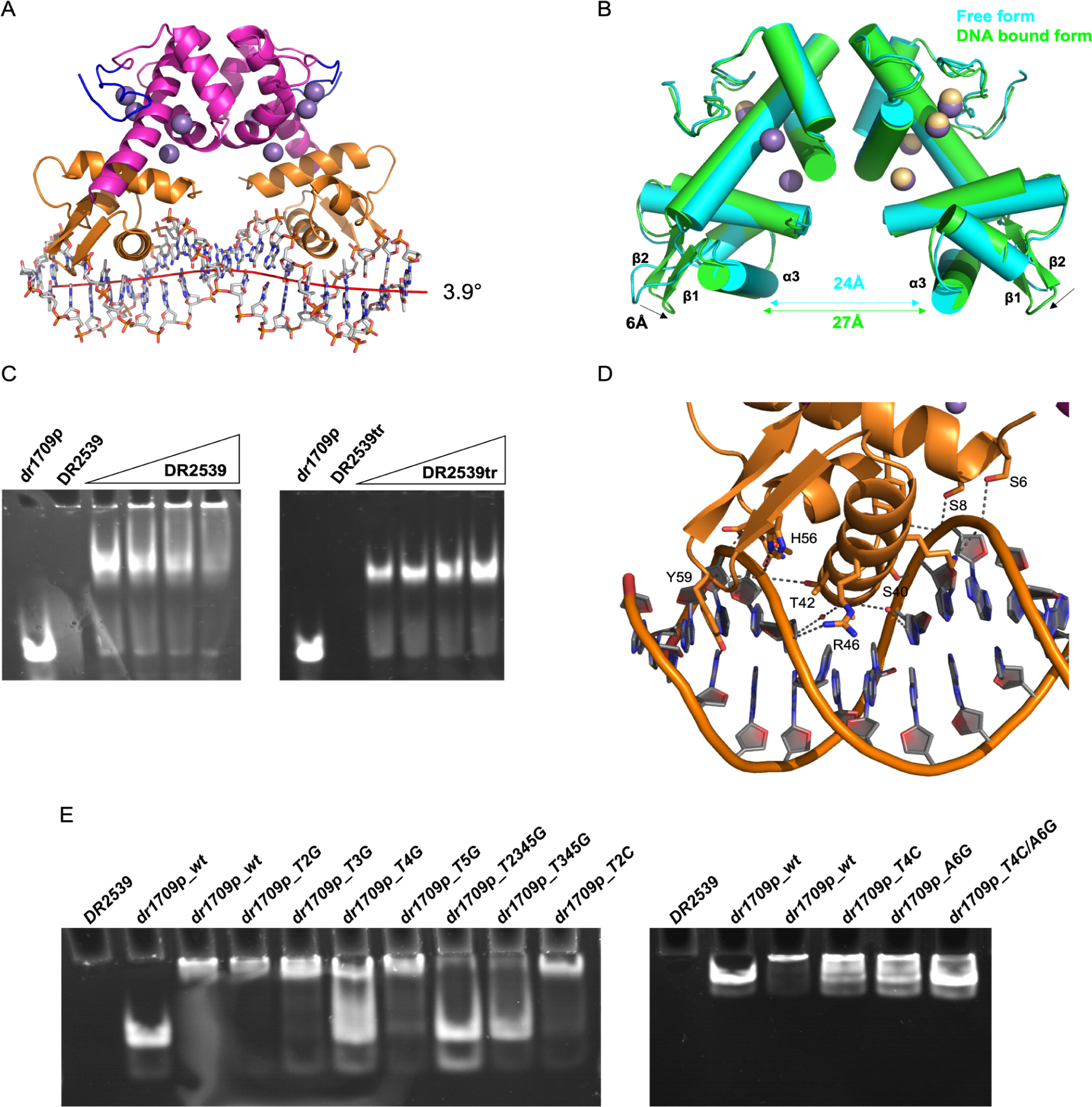
DR2539tr-*dr1709p* interactions. A) Structure of the biological dimer DR2539tr bound to *dr1709* promoter region (21bp) DNA curvature (red line) calculated by CURVES+; B) Superposition of DNA bound (green) and free (green) DR2539tr forms. Manganese and cadmium as purple and wheat spheres, respectively; C) EMSA analysis of *dr1709p* (1µM) binding by titrating the DR2539 (5, 10, 15, 25 µM). On the left, full length DR2539; on the right, truncated DR2539. The non-migration of the band of DR2539 full length at concentrations above 10 µM could be explained either by protein aggregation or formation of bigger complexes. D) Representation of the interactions between one DR2539 monomer (DBD in orange) and the *dr1709 promotor* region. Hydrogen bonds and salt bridges are indicated by dashed grey lines. E) EMSA analysis on the effect of different mutations in the DNA recognition sequence (see Table S5). Left gel, from left to right: DR2539, *dr1709p_wt, dr1709p_wt* and DR2539, *dr1709p_T2G* and DR2539, *dr1709p_T3G* and DR2539, *dr1709p_T4G* and DR2539, *dr1709p_T2345G* and DR2539, *dr1709p_T345G* and DR2539, ND *dr1709p_T2C* and DR2539; Right gel, from left to right: DR2539, *dr1709p_wt, dr1709p_wt* and DR2539, *dr1709p_T4C* and DR2539, *dr1709p_A6G* and DR2539, *dr1709p_T4C/A6C* and DR2539.

The overall structure of DR2539tr in the complex with DNA revealed moderate differences (r.m.s.d. of 1.34 Å) in comparison with free holo-protein, however the DNA binding resulted in an increased distance of 3 Å between carbon-α of the α-helices 3 from both monomers (Figure 2B). DR2539tr dimer binding to DNA B-form results in a slight bending of the DNA, with a total curvature angle of 3.9° (as calculated by CURVES+ [49] (Figure 2A). In fact, the major groove in correspondence of Cyt9 to Gua12 and Thy15 to Ade17, is narrowed by approximately 2 Å compared to canonical B-DNA and accommodates the DNA-binding helices α3 of the HTH motif. Consequently, the minor groove is widened by about 3 Å, and shallower than in a canonical B-form DNA (Figure S5). Circular dichroism spectra titration measurements confirm the DNA bending also in solution, excluding crystallisation artefacts (Figure S6).

### DR2539 recognizes the *dr1709* promotor region

The DR2539 recognizes the *dr1709* promotor region in both full length and truncated forms in presence of manganese (Figure 2C). The region comprising helices α2 and α3 from the HTH motif (residues Ser26-Gln51) located in the DBD, is responsible for most of the DNA binding and recognition (Table S4). Each DBD recognizes a characteristic 5-bases pairs DNA sequence that constitutes the major groove which nests helix α3. However, only a few interactions are occurring in the residue range Ala37-Lys47, (Figure 2D). Several non-specific polar interactions are observed between the DBD and the phosphates of the DNA backbone which are flanking both sides of the major groove. The side chains of Ser26, Thr27, Gln28, Thr42, Arg46, His56 are interacting with the 5’-3’ forward strand and Ser40 and Lys47 with the reverse. The reverse strand also establishes polar interactions with the N-terminal residues, Ser6, Ser8 and Tyr12 (Figure 3). The turn Gly35-Ala39 interacts with nucleotide bases through hydrophobic interactions between Ala37 and Pro38 with C7 methionine groups from reverse thymine16 and forward thymine4, respectively. This interaction defines the specificity for the cognate DNA sequence. Additionally, Ala39 backbone forms hydrogen bonds with two water molecules, and these further interact with O4 from thymine 4 and N4 from reverse cytosine 15.

**Figure 3.**
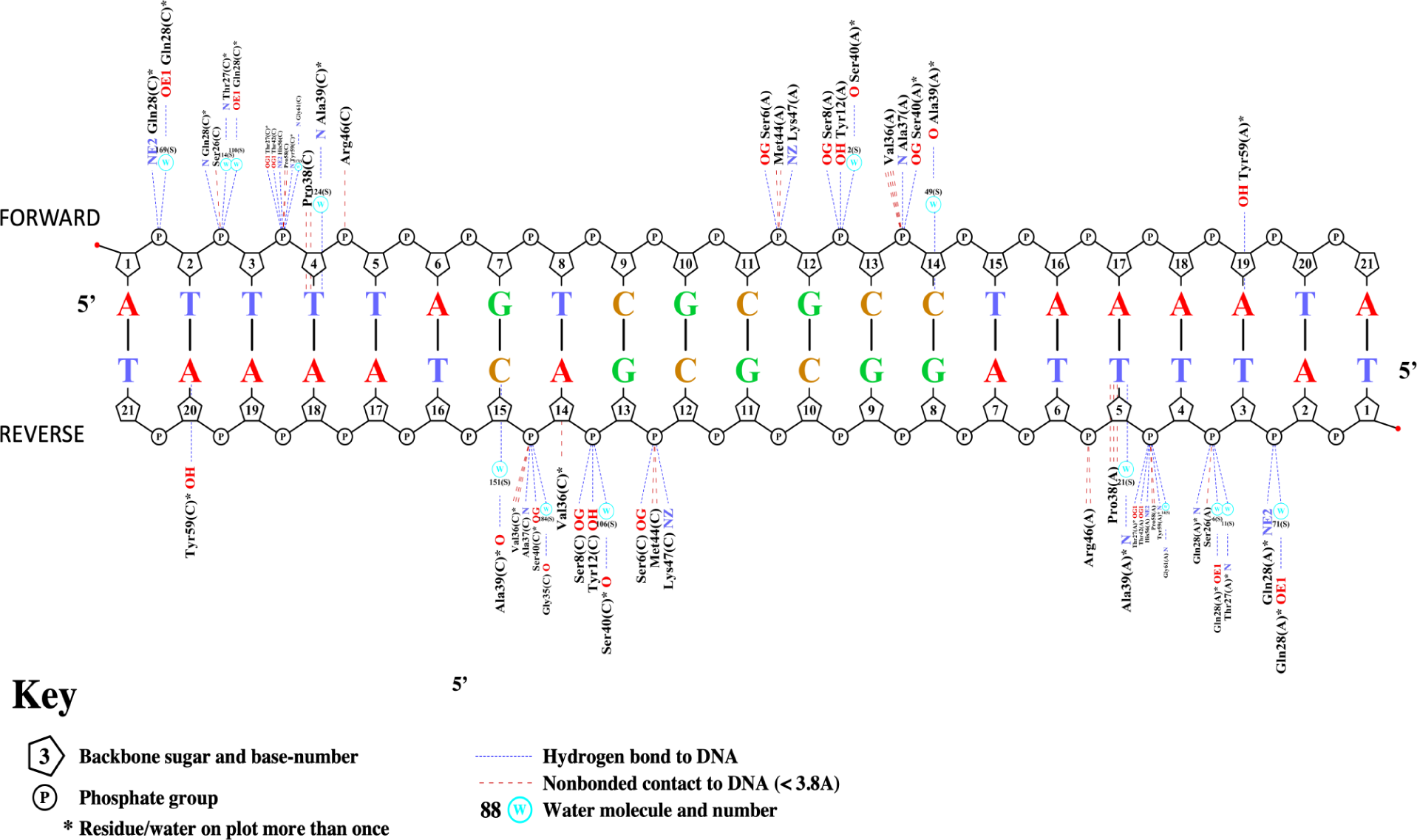
Schematic representation of the specific interactions between DR2539 and *dr1709p*. Figure generated by Nucplot [78].

When with the free form, in the complex, the wings of the winged helix motif, β-strands β1 and β2 from both monomers, are 6 Å closer (distances measured between Tyr59 Cα atoms), pinching the forward strand. Tyr59 is nested into the minor groove and establishes a specific hydrogen bond with the N3 atom from the reverse strand adenine 20 (Figure 2D). Additional interactions are formed between the main chain and DNA mediated by water molecules (Figure 3).

In order to assess the importance of the individual nucleotide bases at different positions, the binding of oligos presenting point mutations on the *dr1709* promotor region (Table S5) to DR2539 was evaluated by gel mobility shift assays (Figure 2E). The hydrogen bond established by Tyr59 and the reverse adenine 20 and the hydrophobic interactions of helix 3 and the major groove were probed in this experiment. The results revealed the role of thymine 4 and reverse thymine16, in which its C7 methyl groups establish hydrophobic interactions with Pro38 and Ala37, respectively. In addition, these two nucleotides share some hydrogen bonded waters with the protein backbone. Mutation of these two nucleotides is enough to dramatically decrease DNA binding (Figure 2E), highlighting the fact that those positions are critical for efficient DNA binding and specificity. We could observe that mutations that affected the hydrophobic interactions (thymine4 and reverse thymine16) were disruptive, while mutations on the reverse adenine 20 and, consequently, hydrogen bond break with the DR2539 wing amino acid, Tyr59, is not essential in the sequence recognition (Figure 2E, but it rather suggests that the role is mainly in the stacking van der Waals interactions of the phenol ring that is nested in the DNA minor groove. Single mutations of forward thymines 3 and 4 showed no shift in the gel while mutation of all nucleotides in the major groove resulted in complete loss of binding (Figure 2E).

### DR2539 presents 4 manganese binding sites

We observed that DR2539 recognizes the *dr1709* promoter region exclusively in the presence of divalent cations. EMSA assays suggest that DR2539tr has affinity for different metal ions, with a distinct preference for transition metals Mn^2+^, Zn^2+^, Fe^2+^, Cd,^2+^ and Ni^2+^ (Figure 4A). The negative control performed without metal supplementation (Figure 4A), indicates that DR2539 can bind to *dr1709p* by uptake of metals either from the media or electrophoresis buffers, while in the presence of EDTA no binding is observed (Figure 4A). The same effect is observed when the DNA is titrated with DR2539 protein and monitored by Circular Dichroism in the 280 nm region (Figure S6). Thus, metal binding is required for the recognition of *dr1709p* DNA sequences by DR2539, providing a direct mechanism connecting metal binding by DR2539 to transcriptional outputs. Contrary to the previously determined structures from the DtxR family, that could bind up to a maximum of three metals [38,45], DR2539 presents four manganese binding sites in each monomer, the first metal binding site we call the primary site (coordinated by Asp11OD1, Glu102OE2, Glu105OE2 and His106NE2), a binuclear ancillary site consisting of ancillary site I and, ancillary site II (coordinated by His79NE2, Glu83OE2, His98ND1, Asp126OD2, His128ND1 and Asp130OD2) and an additional site located in the dimerization domain, here entitled “tertiary site” (coordinated by Glu74OE2, His78NE2, Glu110OE1 and OE2 and Glu113OE1) (Figure 4B). These residues forming the new metal binding site are not conserved in the DtxR family (Figure S2), suggesting that this additional metal site is a peculiarity of DR2539, presumably conserved in *Deinococcus* species [50]. The anomalous Fourier differences map shows strong peaks of 11σ, 9.3σ, 6.5σ and 10.5σ for primary, ancillary sites I and II and tertiary sites, respectively.

**Figure 4.**
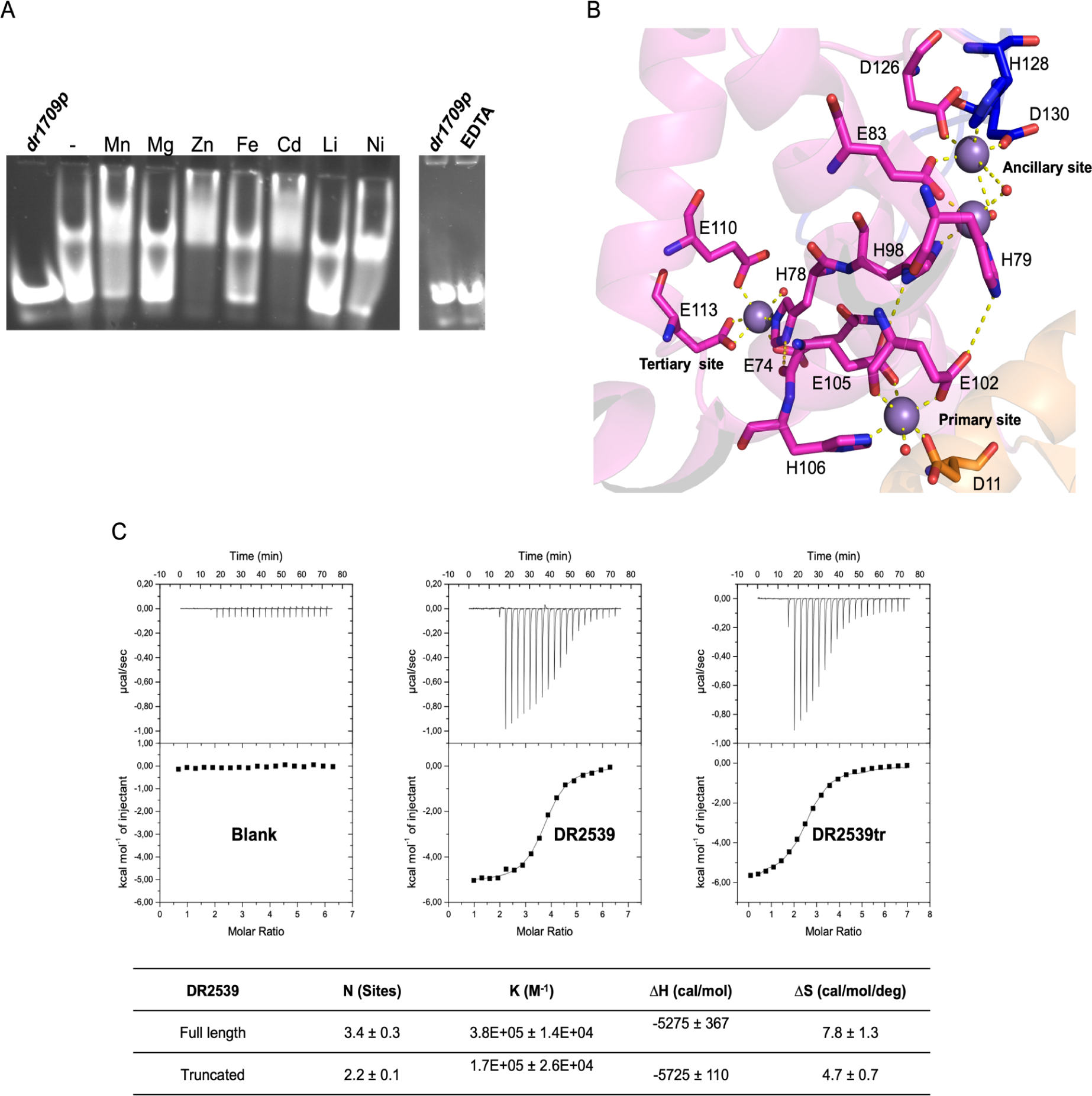
DR2539 binding to metals. A) EMSA analysis of DR2539tr binding to *dr1709p* when incubated with different metal ions (left gel, no metal added (-), manganese (Mn), magnesium (Mg), iron (Fe), cadmium (Cd), lithium (Li), nickel (Ni)) and in the presence of EDTA (right gel). B) The hydrogen bonds network of the manganese binding sites in one of the monomers, manganese ions in purple and water ligands shown as small red spheres. C) Calorimetric titration of DR2539 full length and truncated forms with MnCl_2_, and below the table of thermodynamic parameters calculated by ITC.

Although the incorporation of metals is essential for *in vitro* DR2539 binding to DNA [16], the role of each individual metal-binding site is still unclear. In order to further investigate metal binding, isothermal titration calorimetry (ITC) assays were performed on the full length and truncated versions. The titration curve showed a sigmoidal shape that could be fitted to one set of sites binding model with 3.4 manganese ions (Kd= 2.6 µM) and 2.2 metals (Kd= 5.8 µM) for the full length and the truncated forms, respectively, inferring that the C-terminal domain contributed to the formation of at least one of the binding sites (Figure 4C). The presence of only one transition slope (although at different stoichiometric ratios, for the two forms) indicates that the dissociation constants for the different metal binding sites are in the same order of magnitude.

### Full-length DR2539 presents a compact conformation in solution

The ancillary domain appears to be non-essential for *in vitro* DNA binding, as both truncated and full-length protein presented a similar migration profile in the gel (Figure 4A). Nevertheless, based on the elution profiles on heparin columns, DR2539 full-length and truncated species in presence of manganese ions suggest that they may exhibit different DNA-binding affinities, proposing that the ancillary domain might influence in some manner the DNA binding (Figure S1). Additionally, the ancillary domain has also been associated with protein-protein interaction and the formation of higher complexes, displaying a critical role in vivo [51].

Using AlphaFold2 (AF2) [52], a model for the full-length DR2539 (Figure 5A) was generated and further validated by SAXS experiments. In this model the AD contacts the DD and Asp163 is in position to coordinate the binuclear metal ancillary site (Figure 5B). The position of the ancillary domain is variable in the different crystal structures of the DtxR family and the AD-DD interaction is proposed to be a step of the activation mechanism [47]. In order to probe the interactions of these domains, we performed SAXS experiments. SAXS data revealed that both DR2539 full length and truncated forms are dimers in solution either in apo or holo forms (Table S6), with the calculated Porod’s volumes corresponding to the mass of the dimer. The analysis of the full-length form aimed to identify the position of the ancillary domain and its possible implication in metal regulation. However, the full-length protein was prone to aggregation, especially when incubated with manganese or EDTA (Figure S7). Dilution series were performed to overcome this and the least aggregated samples were obtained for the as-isolated form and the measured scattering curves were analysed. The Guinier region of the scattering curve yields a radius of gyration of 28 Å and analysis of the pair distribution function (P(r)) derived from the scattering curve obtained suggests a maximum intramolecular distance of 83 Å. This is in agreement with the AF2 model since the theoretical scattering calculated for this model reproduces nearly the experimental data with a χ^2^ of 2.9 (Figure 5C). This model also docks into the ab-initio molecular envelope (Figure 5D), indicating that the ancillary domain is likely facing the dimerization domain, allowing it to complete the ancillary metal binding site, likely contributing with a bidentate coordination of Asp163. Furthermore, we wanted to evaluate the possible flexibility of the ancillary domain with respect to the rest of the protein. For that, an ensemble optimization method (EOM) was employed. A pool of 10.000 DR2539 full-length structures was generated, assuming the ancillary domain fully flexible. The minimal ensemble that better fits the experimental data was mainly populated by a conformation with a narrower size distribution than the entire size range of the pool (Figure 5E). This clearly indicates limited flexibility for the AD domain and supports a compacted model in which the ancillary domain should be close to the dimerization domain in the as-isolated form.

**Figure 5.**
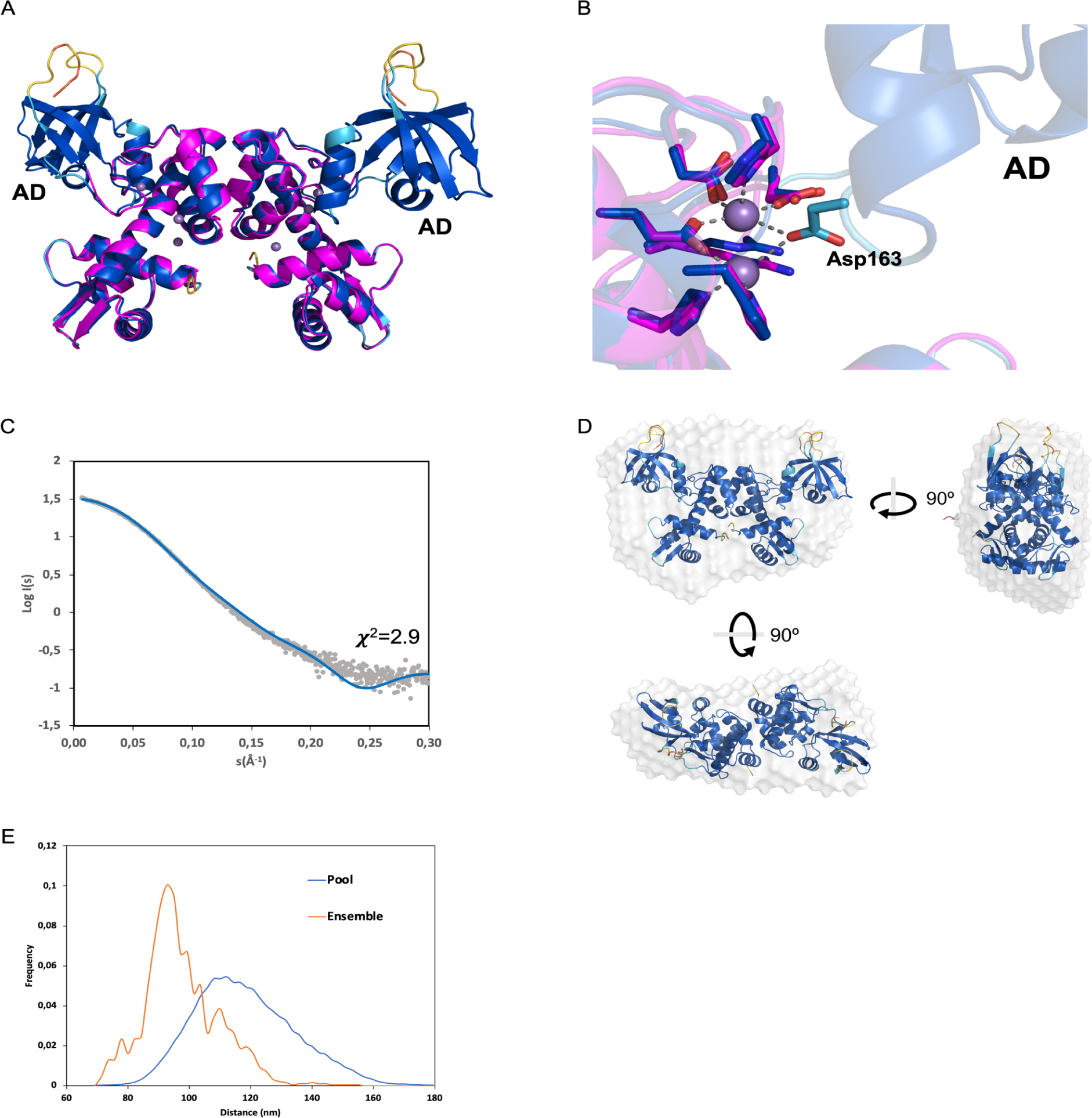
DR2539 full length is a dimer in solution and displays compact arrangement. a) AlphaFold 2 model of full length DR2539 coloured by confidence (dark blue >90% confidence) superposed with experimental structure (PDB ID 8PW0, in magenta). b) Close-up view of the AD-ancillary site interactions with the predicted AF2 model superposed. Manganese ions in purple. c) Experimental SAXS profile by merging the scattering curves obtained at low and high protein concentrations (grey dots) and calculated scattering (blue line) for the DR2539 dimer (model generated by AlphaFold2). d) Ribbon representation of DR2539 model (generated by AF2) docked into the SAXS-derived molecular envelope (semi-transparent surface), generated by DAMMIF, which is the average of 15 independently reconstructed bead models. Three orthogonal views are shown. e) Ensemble optimization method analysis of the flexibility between two rigid segments (DBD-DD and AD) comprising residues 1-126 and 146-233. Frequency distributions of the distances in a pool of models (blue line) and in the selected ensemble that fits the SAXS data of DR2539 (orange line).

As mentioned above, full-length protein was more prone to aggregate in the presence of additives (manganese or EDTA) which made it difficult to analyse SAXS data (Figure S7). Nonetheless, aiming to study the effect of metal binding in the metal regulator activation, we also incubated the as-isolated truncated form with manganese or EDTA. SAXS analysis of the protein solutions in the Guinier region shows a decrease of the Rg upon metal binding, from 27.8 Å in the EDTA incubated sample to 24.6 Å in the manganese incubated sample (Table S6). These results suggest flexibility between DD and DBD and the compact form is achieved at least with insertion of the metal in the primary site. The experimental model (PDB ID 8PVZ) fits better the data of the manganese incubated sample with a χ^2^ of 5.7 (Figure S9).

## Discussion

### The role of DR2539 in *Deinococcus radiodurans* and its structure

Metalloregulators are key controllers in maintaining bacterial metal homeostasis by sensing the cellular metal abundance or deficiency. This work highlights the structural and biophysical features of the manganese homeostasis regulator, DR2539, crucial in the cellular oxidation response of *Dr* [16]. The overall structure of DR2539 presents the three characteristic domains of the DtxR family: the N-terminal DNA-binding domain (DBD), the dimerization domain and the ancillary domain. The latter is absent in our crystal structure due to spontaneous proteolysis. Although the overall fold is conserved, this protein revealed unique features both in metal incorporation and in protein-DNA interactions that could be related to the uncommon Mn homeostasis of *Dr*. Unlike other reported members of the DtxR family, for which it was proposed that the metal cofactor binds to the monomer and consequently promoting conformational changes that induce the dimerization, our study reveals dimerisation is not metal dependent, with no evidence for monomeric DR2539 in the absence of metals [46,47].

### DR2539 metal binding sites in comparison with other DtxR family members

DR2539 presents structural and functional differences from *Cd*DtxR homologues. The structure of the ternary complex (DR2539tr:DNA:Mn) unveils four manganese binding sites. The primary binding site in DR2539 is constituted by acidic residues, characteristic of manganese specific regulators, whereas iron specific binding regulators present sulphur containing amino acids. Contrary to the *Bs*MntR and *Bh*MntR homologues, which present a binuclear centre, with a glutamate coordinating the second metal in a bidentate fashion, DR2539 presents Lys14 in this position, which impairs the formation of a binuclear centre in the DBD-DD interface. The presence of a positive charge is also observed in *Cd*DtxR and *Mt*IdeR with an arginine at this position. On the other hand, DR2539 presents a bimetal centre in the ancillary domain, which further contributes to anchoring this domain and modulating the activity of the repressor. The ancillary site, that in the case of *Cd*DtxR and *Mt*IdeR is a mononuclear centre and it is coordinated by two residues from the ancillary domain, in DR2539, *Sm*SloR, *Sg*Scar and *Mt*MntR, it is coordinated by only one residue from the AD, Asp163 (DR2539 numbering). Moreover, although the second metal is not directly coordinated to the AD, it aids the interaction between DD and AD. While in DR2539 this bimetal site is shaped by three histidines (His79, His98 and His128), three aspartates (Asp126, Asp130 and Asp163) and one Glu83, *Sm*Slor, *Sg*ScaR, *Sp*MtsR and *Ta*IdeR present a cysteine replacing the first aspartate residue (Figure S2). The presence of binuclear centres appears to be a peculiarity of Mn activated regulators from DtxR family, at the point that it has been suggested that the geometry of the binuclear centre serves to select manganese over other transition metals, which is likely the case, given the versatility of histidine in binding metals [53].

The new metal binding site identified in the dimerization domain, between helices α4 and α6, suggests a role in stabilising the interaction between the dimerization and DNA-binding domains. The presence of a fourth metal in the tertiary site, is confirmed by stoichiometric binding ratio of 3.4 manganese ions to the full-length protein (Figure 4C) and it seems to be a peculiarity of DR2539. The four metal binding sites are part of a hydrogen bonding network, suggesting that, despite the similar affinities, the metal incorporation is part of a step-by-step mechanism. In the holo-form, Glu102 hydrogen bonds to His98 and Glu105 to His79 (Figure 2D). Furthermore, His78, coordinating the tertiary site, is only 3.8 Å from the backbone carbonyl of Glu105, which coordinates the primary site.

According to the proposed *Cd*DtxR mechanism, the ancillary site is the highest affinity site, and the primary site is the low affinity site [47]. However, considering manganese anomalous signals in the ternary complex, we found that the primary and tertiary sites show higher anomalous differences, pointing to tighter binding compared to the ancillary sites. We should nevertheless consider the fact that the ancillary domain is missing, hence the corresponding metal binding site could be loose due to the lack of the coordinating residue (Asp163). Furthermore, in solution, isothermal titration analysis showed that the truncated form loses one metal compared to the full-length protein. This result, supported by the full length AlphaFold model, suggests that the ancillary domain may be involved in the coordination of the missing metal.

### *dr1709p* sequence recognition by DR2539

According to the mechanism proposed for *Cd*DtxR [47,54], a multistep metal binding induces conformational changes that promote protein-DNA interaction. The N-terminal amino acids Ser6, Ser8 and the conserved Tyr12 establish hydrogen bonds with DNA phosphate groups promoted by the conformation held by the primary metal site, while it presents a limited number of salt bridge interactions (only Arg46 and Lys47 from α3 helix) when compared with the structures of *Mt*IdeR and *Cd*DtxR in complex with DNA [55] that show an extended positively charged patch that interacts with the negatively charged phosphate backbone.

Helix α3, that is nested in the DNA major groove, reveals a close sequence similarity among groups of subfamilies: *Cd*DtxR, *Se*IdeR and *Mt*IdeR; *Sg*ScaR, *Sm*SloR, and *Sp*MtsR; DR2539 and *Mt*MntR; and *Ta*IdeR (Figures S2 and S8). In the first two subfamilies, the helix α3 first residue is a proline that participates in the interface with the DNA. In DR2539, the first residue of α3 is Ala39, which interacts directly with Thy5 of the forward filament and Thy16 of the reverse (Figure 3). In this position a bulkier side chain would reshape the interface, likely diminishing the affinity for the DNA strand. *Mt*MntR does not present any proline in the region, and the first residue is a serine suggesting a different protein-DNA interaction [38].

The fifth residue of helix α3 in the homologous proteins (Gly43 in DR2539, Figure S2) has been reported [56,57] to interact directly by hydrogen bonding with the nucleotides of a G-C base-pair and to recognise the correct DNA sequence [57]. While homologous proteins have a polar residue with long side chain, DR2539 presents instead Gly43, which permits to recognise and bind either binding boxes of *dr1709p*, and the variability in the DNA sequence in the position #8, that breaks the palindrome sequence.

### DR2539 regulation mechanism

DR2539 is a dimer in the apo-form and is fully loaded with 8 manganese ions in the holo-form. Primary and ancillary sites are conserved from homologous proteins, but an additional metal binding site in the dimerization domain was observed for the first time, contributing to increase metal:protein activation ratio.

The flexibility between DBD and DD suggested that metal binding site 1 is critical to DR2539 scaffold and DNA recognition, with SAXS data supporting that manganese ion binding on the primary site is responsible for the stabilisation of the DBDs and the correct spacing between them. The DBD-DNA interaction presents unique features, differently from the N-terminal helix-to-coil transition induced by metal binding and DNA-induced conformational changes observed in *Cd*DtxR [57,58]. In DR2539, the N-terminal residues are in the opposite direction compared to *Cd*DtxR:DNA complex (PDB ID: 1C0W) and Ser6 establishes a hydrogen bond with O from phosphate of DNA backbone. Additionally, contrary to *Cd*DtxR, the N-terminal is not making any interaction with the primary binding site. Binding of DNA to DR2539 holo-form triggered conformational changes of the DBD orientation, thereby favouring the interaction of certain residues with DNA (Figure 3). The distance between the α3 recognition helices is increased to match the spacing between two consecutive major groove regions. Additionally, Tyr59 from the antiparallel β-sheet motifs, interacts with adenines 19 and 20 from the forward and reverse strands, respectively. Although the tyrosine may play a role in a latch closing mode which is usually occupied by charged residues in the homologous proteins, the perturbation of the hydrogen bond caused no visible effects in the “*in vitro*” qualitative binding experiments (Figure 2E). On the other hand, the recognition of the major groove is essential with special attention to interactions to thymine 4 and reverse thymine 16 that are crucial in recognition, although single mutations do not completely impair “*in vitro*” DNA binding.

### The role of the ancillary domain in DR2539

In *Cd*DtxR and *Mt*IdeR, the C-terminal domain is either coordinating the ancillary metal site and interacting with dimerization and DNA binding domains or in variable positions, stabilised by crystallographic contacts [55,59]. Although several structures the DtxR family lack electron density to model this domain due to the flexibility provided by the long linker, most of the DR2539 linker (Pro127-Pro140) is well defined in the electron density map of the truncated protein and forms a double hairpin, with the first turn involved in the formation of the ancillary metal binding site. The long linker connecting the dimerization and ancillary domains is compatible with an extended apo-form in solution, however the EOM analysis on the SAXS data excludes a high degree of flexibility of this region, suggesting that, in the as-isolated form, the AD is close to the DD in a compact form. Although, in absence of metals, DR2539 presented a tendency to aggregate, potentially indicating an important change in protein globularity.

The structural comparison with *Sg*ScaR, *Sm*SloR and *Mt*MntR and the AlphaFold model reveals the presence of structurally conserved aspartate residues (Asp163 in DR2539) that participate in forming the ancillary metal binding site and could contribute to structure the double metal binding site, modulating the repressor activation in function of the metal ion concentration.

Additional sequence analysis identifies this domain as a member of the FeoA superfamily (pfam 04023). FeoA is a small cytoplasmic beta-barrel protein that presents a SH3-like fold (NMR structural studies of E. coli FeoA (PDB ID:2LX9). SH3-like domains are usually involved in protein-protein interactions [60]. Although protein oligomerization through the C-terminal domain was proposed for the *Sp*MstR and *Se*IdeR homologues [51][61], it is not consensual that this behaviour is common in the entire DtxR family. The amino acids proposed in protein-protein interactions (Y167 and F187 in *Sp*MtsR to T171 and A191 in DR2539, respectively) are not conserved in DR2539, however their chemical nature is conserved (Figure S2), therefore, it cannot be excluded a similar oligomerization behaviour in compatible conditions.

Although the ancillary domain is not essential for in vitro DNA binding, we observe some differences between full length and truncated forms. The AD modulates the binding, probably through the metal incorporation into the ancillary metal binding site, which stabilises the entire scaffold. These results agree with *Cd*DtxR *in vivo* studies, showing that the absence of the ancillary domain leads to a less active repressor [54].

## Conclusion

The current study elucidates the structural determinants of the metal regulator DR2539 in metal binding and DNA interaction. This regulator is revealed to be key in manganese homeostasis and consequently in ROS scavenging. DR2539 exhibits specificity for manganese binding enabling the transcription factor to bind DNA and activate its target genes, including the known manganese transporter *dr1709*. When compared to homologues, DR2539 presents the highest metal loading capacity, which, along with lower protein-*dr1709p* specific interactions, may contribute to enhancing concentration of manganese in *Dr*. The presence of the ancillary domain, while it is not essential for *in vitro* DNA regulation, shapes the binuclear ancillary metal binding site, modulating the DNA binding. Although no higher oligomeric forms were observed in this study, we cannot exclude that *in vivo* additional cofactors or higher oligomerisation states may be present due to assembly of multiple factors on the genomic DNA.

## Material and methods

### Cloning, expression and purification

The gene encoding DR2539 protein was amplified by polymerase chain reaction using the genomic DNA of *Deinococcus radiodurans* as a template. It was inserted into the *EheI/HincIII*-digested expression vector pProEx HTb (Life Technologies), resulting in a 25-residues hexahistidine-containing tag at the N-terminus. The recombinant DR2539 was expressed in *Escherichia coli* BL21 (DE3) (Invitrogen). The cells were grown at 310 K to an OD600 of ≈ 0.6 in Luria-Bertani medium containing 100 µg ml^-1^ of ampicillin. Protein expression was induced using 1 mM isopropyl β-D-1-thiogalactopyranoside (IPTG). Cell growth continued at 289 K for 14 h after IPTG induction and the cells were harvested by centrifugation at 7000 g for 20 min at 277 K.

The cell pellet was resuspended in lysis buffer (50 mM Tris-HCl pH 7.5, 1M NaCl, 10% (v/v) glycerol, 2 mM β-mercaptoethanol, 1.5 M urea) to which DNAse I was added to a final concentration of 20 µg/ml together with an EDTA-free protease inhibitor tablet (Roche Applied Science). Cells were disrupted by 3 cycles of freeze/thaw and the suspension was then homogenized using an ultrasonic processor (QSonica). The crude cell extract was centrifuged at 40 000 g for 45 min at 277 K. The supernatant was filtered in 0.45 µm filter and loaded into a 5 ml His-trap FF column (GE Healthcare) pre-equilibrated with buffer A (50 mM Tris-HCl pH 7.5, 500 mM NaCl, 5% (v/v) glycerol, 1.5 M urea, 5 mM imidazole). The column was washed with buffer B (50 mM Tris-HCl pH 7.5, 500 mM NaCl, 5% (v/v) glycerol, 5 mM imidazole). A first washing step was performed with 10% buffer C (50 mM Tris-HCl pH 7.5, 500 mM NaCl, 5% (v/v) glycerol, 500 mM imidazole) followed by a gradient from 10% to 100% of buffer C, in 20 column volumes. The eluted fractions purity were evaluated by SDS-PAGE, with two forms of DR2539 (truncated and full-length) being eluted around 250 mM imidazole. Protein fractions were pooled and tobacco etch virus (TEV) protease (ratio 1:20 (w/w) added. The buffer was exchanged to TEV cleavage buffer D (50 mM Tris-HCl pH7.5, 500 mM NaCl, 5% (v/v) glycerol, 1 mM DTT) in an overnight dialysis at 277 K. After incubation, the protein mixture was loaded into a 5 ml His-trap FF column (GE Healthcare) pre-equilibrated with buffer B. The mixture of protein truncated and full length was eluted with a elution step of 30% buffer C and the buffer exchanged to buffer E (50 mM Tris-HCl pH 7.5, 150 mM NaCl, 1mM MnCl_2_) using a centricon concentrator with a cutoff of 10 kDa (Millipore). The sample was applied onto a Heparin column (GE Healthcare) which had previously been equilibrated with buffer E. The two forms (full-length and truncated) were separated with a linear gradient up to 100% of buffer F (50 mM Tris-HCl pH 7.5, 1M NaCl, 1mM MnCl_2_) over 8 column volumes. The last step was a gel filtration on a Superdex 75 column (GE Healthcare) equilibrated with 50 mM Tris-HCl pH 7.5, 300 mM NaCl. The purified samples were concentrated to 10 mg ml^-1^, flash-frozen in liquid nitrogen and stored at 193 K until further use.

### Preparation of double-stranded DNA

Forward and reverse DNA oligonucleotides containing the different sequences designed for DNA-binding analysis and crystallization studies were obtained from Eurofins (Table S5). To prepare double-stranded DNA, each forward and reverse oligonucleotide sample pair was resuspended in 10 mM Tris-HCl pH 7.5 and 100 mM NaCl and mixed to a final concentration of 100 µM for EMSA and 1M for crystallization assays. All mixtures were then heated at 95°C for 5 min, slowly cooled down to 56°C for 30 min in a thermal cycler and left for 1h at room temperature.

### Crystallization, data collection, structure solution and refinement

Initial crystallization screenings were performed by sitting-drop vapour diffusion method at 277 and 293 K using Greiner CrystalQuick 96 well plates. A Cartesian PixSys 4200 crystallization robot (Genomic solutions, UK) facility (High Throughput Crystallization Laboratory (HTX Lab)) at EMBL Grenoble) was used to dispense 200 nl drops with 1:1 protein:precipitant ratio.

DR2539 truncated protein was initially screened by the following commercial crystallization screens: Classics Suite (Qiagen), Salt grid (Hampton Research), Wizard I&II (Rigaku reagents) and JCSG + (Molecular Dimensions). A cluster of needles appeared within 15 days on condition D3 from the Classics suite plate (0.1 M Hepes pH 7.5; 1 M sodium acetate; 0.05 M Cadmium sulfate). In order to improve crystal quality and size, the condition was scaled up to 2 μl drops with 1:1 protein:precipitant ratio, and precipitant concentration and pH were screened. Single crystals were obtained after 3 days in sitting drops (0.1 M Hepes pH 7.0; 1.2 M sodium acetate; 0.05 M Cadmium sulphate) and grew up to a maximum size of 0.02 x 0.02 x 0.4 mm in 7 days. An additive screen (Hampton Research) was performed and better diffracting crystals were obtained in condition C6 (3% 6-Aminohexanoic acid; 0.1M Hepes pH 7.0; 1.2 M sodium acetate; 0.05M Cadmium sulphate).

To prepare the complex, Dr2539tr was incubated with *dr1709p_wt* and manganese (410 µM, 205 µM and 820 µM, respectively) in 20 mM Tris-HCl pH7.5, 150 mM NaCl. The complex was initially screened by the following commercial crystallization screens: Nucleix (Qiagen) and Wizard I&II (Rigaku reagents). Single crystals were obtained in condition F12 (PEG 8K 30% (w/v); Imidazole 0.1 M pH 8.0; NaCl 0.2 M) within 24 h and diffracted up to 4 Å. The condition was scaled up to 2 μl hanging drops with 1:1 protein-promotor precipitant ratio, and the precipitant concentration and pH were screened. Single plates were obtained within 24h (PEG 8K 26% (w/v); Imidazol 0.1 M pH 7.8; NaCl 0.2 M) and grew up to a maximum size of 0.05 x 0.02 x 0.2 mm in 7 days. Crystals were transferred into a cryoprotectant solution consisting of a reservoir solution augmented with 20% (v/v) glycerol and then flash-cooled in liquid nitrogen. X-ray diffraction data sets were collected at 100 K using a Pilatus 6M-F detector on ID29 [62] and ID23-1 [63] beamlines at the European Synchrotron Radiation Facility. A highly redundant anomalous dispersion data set was collected 1.771 Å on ID29 in order to obtain the maximum anomalous signal from cadmium (theoretical K-edge absorption peak at (3.085 Å)) without compromising the beam intensity. X-ray diffraction images were integrated and scaled with XDS [64]. The SHELX [65] package was used to estimate the amount of anomalous signal, find the positions of the cadmium atoms and calculate initial phases. ARP/wARP [66] was used to build an initial model with the docked sequence. Finally, manual model building and interactive refinement were performed using Coot [67] and PHENIX [68], which was also used to refine the metal occupancies.

Final model was refined with a higher resolution dataset collected from a native crystal at 0.980 Å on ID23-1. X-ray diffraction data set from a crystal of DR2539 in complex with *dr1709* promotor was collected on ID29. The structure was solved by molecular replacement with Phaser [69] using the structure of DR2539 truncated obtained before as the search model. Model building and refinement were performed with Coot and PHENIX.

### Small-angle X-ray Scattering

Small-angle X-ray Scattering (SAXS) measurements were carried out at 293 K on beamline BM29 [70] of the ESRF at λ =0.9919 Å. Scattering curves were recorded over a scattering-vector of 0.0035<q<0.5Å^-1^, (q = 4π sin (θ)/λ, 2θ is the scattering angle) using a Pilatus 1M detector (Dectris Ltd., Baden, Switzerland). Prior to experiments all solutions were centrifuged at 12,000 x g for 10 minutes to remove aggregated particles. 50 μL of protein solution, loaded in a flow mode into a sample capillary using a liquid handling robot, were exposed to X-rays and scattering data collected using multiple exposures (10 exposures each of 10 seconds). Matched buffer measurements taken before and after every sample measurement were averaged and used for background subtraction. Data were processed and analysed with the ATSAS package [71,72]. Merging of scattering curves obtained at different sample concentrations and downstream analysis were performed manually as described in the literature [73] in order to obtain the zero concentration curve.

Forward scattering I(0) and radius of gyration (Rg) were calculated from the Guinier approximation [74]. Particle volume and the maximum particle size (Dmax) were determined from the pair distribution function P(r) as computed by GNOM [75] using PRIMUS [73]. Scattering curves from solutions of full-length and truncated forms were measured at different concentrations in buffer 50 mM Tris-HCl pH 7.5, 300 mM NaCl. The proteins were also dialysed in buffer supplemented with 5 mM EDTA or 5 mM MnCl2. *Ab* initio models of the solution shape of DR2539 were derived from the experimental scattering curve using DAMMIF [76] imposing P2 symmetry. To produce a final molecular envelope 15 independently generated *ab initio* models were aligned, averaged and filtered using DAMAVER [77]. Theoretical scattering curves of DR2539 AlphaFold2 model [52], or the experimental model (PDB ID 8PVZ), were calculated and used to generate fits against experimentally-obtained scattering curves using CRYSOL [71].

### Electrophoretic mobility shift assay

The electrophoretic mobility shift assays (EMSAs) were performed as previously described [16] with some modifications. 5-25 μM of DR2539 (full-length or truncated) was incubated for 30 min at 4°C with 1 µM of the annealed oligos (Table S5) in EMSA binding buffer (25 mM Tris-HCl pH7.5, 150 mM NaCl, 0.5 mM MnCl_2_ and 5% glycerol) to a final volume of 20 µl. Different salts were tested (MnCl_2_, MgCl_2_, ZnCl_2_, (NH_4_)_2_Fe(SO_4_), CdSO_4_, LiCl, NiCl_2_) at 0.5 mM and EDTA at 1 mΜ. After the incubation, the reaction mixtures were analysed on 5% nondenaturing polyacrylamide gels. The gels were incubated with SYBR green (Invitrogen) and revealed under an UV transilluminator.

### Isothermal Calorimetry

Isothermal calorimetry assays were conducted at 298 K on a Microcal ITC 200 (Microcal Inc., Northampton, MA, USA) with the reference cell filled with MilliQ water. Proteins were treated with EDTA (10 mM) in the storing buffer (25 mM Tris-HCL pH 7.5, 300 mM NaCl) and extensively dialysed against the titration buffer (50 mM Tris-HCl pH7.5, 300 mM NaCl). Full-length and truncated forms of DR2539 at 50 μM were titrated with 20 injections of 2 uL of MnCl_2_ (1.5 mM). After subtraction of the baselines, the integrated heat responses were fitted using Origin software package (Northampton, MA, USA).

### Circular Dichroism Measurements

Binding of DR2539 to *dr1709p* was monitored by circular dichroism (CD) spectroscopy. CD spectra (260-320 nm) were recorded at 293 K in 1-nm steps on a temperature controlled Jasco J-815 spectropolarimeter, continuously purged with nitrogen gas. For each sample the smoothed average of 16 spectra was considered. DR2539 protein was added incrementally to a 10 mm-path quartz cuvette (500 μL) containing *dr1709p* (2 μM) in binding buffer (25 mM Tris-HCl pH7.5, 150 mM NaCl, 0.5 mM MnCl2) and metal depletion test was performed in buffer supplemented with 2 mM EDTA.

## Acknowledgments

CM acknowledges the ‘Fundação para a Ciência e Tecnologia’ (FCT) for a postdoctoral contract SFRH/BEST/51724/2011. DdS and CM acknowledge Programa Pessoa from FCT and Campus France ref. 35797UH. The authors also thank the Partnership for Structural Biology, Grenoble (France) for an integrated structural biology environment and the ESRF Structural Biology Group for continuous support. Salette Reis and Claudia Pinho are gratefully acknowledged for assistance with the ITC experiments. Pedro Pereira is gratefully acknowledged for helpful discussions on the project.

## References

1 Slade D & Radman M (2011) Oxidative stress resistance in Deinococcus radiodurans. Microbiol Mol Biol Rev 75, 133–191.

2 Anderson A, Nordon H, Cain RF, Parrish G, Duggan D, Anderson A, Nordan H, Parish G & Cullum-Dugan D (1956) Studies on a radio-resistant micrococcus. I. Isolation, morphology, cultural characteristics, and resistance to gamma radiation. Food Technol.

3 Driedger AA (1970) The DNA content of single cells of Micrococcus radiodurans. Can J Microbiol 16, 1136–1137.

4 Cox MM (1999) Recombinational DNA repair in bacteria and the RecA protein. Prog Nucleic Acid Res Mol Biol 63, 311–366.

5 Cox MM, Goodman MF, Kreuzer KN, Sherratt DJ, Sandler SJ & Marians KJ (2000) The importance of repairing stalled replication forks. Nature 404, 37–41.

6 Kowalczykowski SC (2000) Initiation of genetic recombination and recombination-dependent replication. Trends Biochem Sci 25, 156–165.

7 Kuzminov A (2001) DNA replication meets genetic exchange: chromosomal damage and its repair by homologous recombination. Proc Natl Acad Sci U S A 98, 8461–8468.

8 Daly MJ, Gaidamakova EK, Matrosova VY, Vasilenko A, Zhai M, Leapman RD, Lai B, Ravel B, Li S-MW, Kemner KM & Fredrickson JK (2007) Protein oxidation implicated as the primary determinant of bacterial radioresistance. PLoS Biol 5, e92.

9 Daly MJ (2009) A new perspective on radiation resistance based on Deinococcus radiodurans. Nat Rev Microbiol 7, 237–245.

10 Makarova KS, Aravind L, Daly MJ & Koonin EV (2000) Specific expansion of protein families in the radioresistant bacterium Deinococcus radiodurans. Genetica 108, 25–34.

11 Daly MJ, Gaidamakova EK, Matrosova VY, Kiang JG, Fukumoto R, Lee D-Y, Wehr NB, Viteri GA, Berlett BS & Levine RL (2010) Small-molecule antioxidant proteome-shields in Deinococcus radiodurans. PLoS One 5, e12570.

12 Sun H, Xu G, Zhan H, Chen H, Sun Z, Tian B & Hua Y (2010) Identification and evaluation of the role of the manganese efflux protein in Deinococcus radiodurans. BMC Microbiol 10, 319.

13 Makarova KS, Omelchenko MV, Gaidamakova EK, Matrosova VY, Vasilenko A, Zhai M, Lapidus A, Copeland A, Kim E, Land M, Mavrommatis K, Pitluck S, Richardson PM, Detter C, Brettin T, Saunders E, Lai B, Ravel B, Kemner KM, Wolf YI, Sorokin A, Gerasimova AV, Gelfand MS, Fredrickson JK, Koonin EV & Daly MJ (2007) Deinococcus geothermalis: the pool of extreme radiation resistance genes shrinks. PLoS One 2, e955.

14 Chang S, Shu H, Li Z, Wang Y, Chen L, Hua Y & Qin G (2009) Disruption of manganese ions [Mn(II)] transporter genes DR1709 or DR2523 in extremely radio-resistant bacterium Deinococcus radiodurans. Wei Sheng Wu Xue Bao 49, 438–444.

15 Bozzi AT, Bane LB, Weihofen WA, Singharoy A, Guillen ER, Ploegh HL, Schulten K & Gaudet R (2016) Crystal Structure and Conformational Change Mechanism of a Bacterial Nramp-Family Divalent Metal Transporter. Structure 24, 2102–2114.

16 Sun H, Li M, Xu G, Chen H, Jiao J, Tian B, Wang L & Hua Y (2012) Regulation of MntH by a dual Mn(II)- and Fe(II)-dependent transcriptional repressor (DR2539) in Deinococcus radiodurans. PLoS One 7, e35057.

17 Chen H, Wu R, Xu G, Fang X, Qiu X, Guo H, Tian B & Hua Y (2010) DR2539 is a novel DtxR-like regulator of Mn/Fe ion homeostasis and antioxidant enzyme in Deinococcus radiodurans. Biochem Biophys Res Commun 396, 413–418.

18 Santos SP, Mitchell EP, Franquelim HG, Castanho MARB, Abreu IA & Romão CV (2015) Dps from Deinococcus radiodurans: oligomeric forms of Dps1 with distinct cellular functions and Dps2 involved in metal storage. FEBS J 282, 4307–4327.

19 Chen H, Xu G, Zhao Y, Tian B, Lu H, Yu X, Xu Z, Ying N, Hu S & Hua Y (2008) A novel OxyR sensor and regulator of hydrogen peroxide stress with one cysteine residue in Deinococcus radiodurans. PLoS One 3, e1602.

20 Ul Hussain Shah AM, Zhao Y, Wang Y, Yan G, Zhang Q, Wang L, Tian B, Chen H & Hua Y (2014) A Mur regulator protein in the extremophilic bacterium Deinococcus radiodurans. PLoS One 9, e106341.

21 Haemig HAH, Moen PJ & Brooker RJ (2010) Evidence that Highly Conserved Residues of Transmembrane Segment 6 of Escherichia coli MntH Are Important for Transport Activity. Biochemistry 49, 4662.

22 Gunshin H, Mackenzie B, Berger UV, Gunshin Y, Romero MF, Boron WF, Nussberger S, Gollan JL & Hediger MA (1997) Cloning and characterization of a mammalian proton-coupled metal-ion transporter. Nature 388, 482–488.

23 Anderson ES, Paulley JT, Gaines JM, Valderas MW, Martin DW, Menscher E, Brown TD, Burns CS & Roop RM 2nd (2009) The manganese transporter MntH is a critical virulence determinant for Brucella abortus 2308 in experimentally infected mice. Infect Immun 77, 3466–3474.

24 Cellier M, Privé G, Belouchi A, Kwan T, Rodrigues V, Chia W & Gros P (1995) Nramp defines a family of membrane proteins. Proc Natl Acad Sci U S A 92, 10089–10093.

25 Makui H, Roig E, Cole ST, Helmann JD, Gros P & Cellier MF (2000) Identification of the Escherichia coli K-12 Nramp orthologue (MntH) as a selective divalent metal ion transporter. Mol Microbiol 35, 1065–1078.

26 Courville P, Chaloupka R & Cellier MFM (2006) Recent progress in structure-function analyses of Nramp proton-dependent metal-ion transporters. Biochem Cell Biol 84, 960–978.

27 Boyd J, Oza MN & Murphy JR (1990) Molecular cloning and DNA sequence analysis of a diphtheria tox iron-dependent regulatory element (dtxR) from Corynebacterium diphtheriae. Proc Natl Acad Sci U S A 87, 5968–5972.

28 Feese MD, Ingason BP, Goranson-Siekierke J, Holmes RK & Hol WG (2001) Crystal structure of the iron-dependent regulator from Mycobacterium tuberculosis at 2.0-A resolution reveals the Src homology domain 3-like fold and metal binding function of the third domain. J Biol Chem 276, 5959–5966.

29 Manabe YC, Saviola BJ, Sun L, Murphy JR & Bishai WR (1999) Attenuation of virulence in Mycobacterium tuberculosis expressing a constitutively active iron repressor. Proc Natl Acad Sci U S A 96, 12844–12848.

30 Yeo HK, Park YW & Lee JY (2014) Structural analysis and insight into metal-ion activation of the iron-dependent regulator from Thermoplasma acidophilum. Acta Crystallogr D Biol Crystallogr 70, 1281–1288.

31 Marcos-Torres FJ, Maurer D, Juniar L & Griese JJ (2021) The bacterial iron sensor IdeR recognizes its DNA targets by indirect readout. Nucleic Acids Res 49, 10120–10135.

32 Stoll KE, Draper WE, Kliegman JI, Golynskiy MV, Brew-Appiah RAT, Phillips RK, Brown HK, Breyer WA, Jakubovics NS, Jenkinson HF, Brennan RG, Cohen SM & Glasfeld A (2009) Characterization and structure of the manganese-responsive transcriptional regulator ScaR. Biochemistry 48, 10308–10320.

33 Glasfeld A, Guedon E, Helmann JD & Brennan RG (2003) Structure of the manganese-bound manganese transport regulator of Bacillus subtilis. Nat Struct Biol 10, 652–657.

34 Ando M, Manabe YC, Converse PJ, Miyazaki E, Harrison R, Murphy JR & Bishai WR (2003) Characterization of the Role of the Divalent Metal Ion-Dependent Transcriptional Repressor MntR in the Virulence of Staphylococcus aureus. Infect Immun 71, 2584.

35 Horsburgh MJ, Wharton SJ, Cox AG, Ingham E, Peacock S & Foster SJ (2002) MntR modulates expression of the PerR regulon and superoxide resistance in Staphylococcus aureus through control of manganese uptake. Mol Microbiol 44, 1269–1286.

36 Que Q & Helmann JD (2000) Manganese homeostasis in Bacillus subtilis is regulated by MntR, a bifunctional regulator related to the diphtheria toxin repressor family of proteins. Mol Microbiol 35, 1454–1468.

37 Lee MY, Lee DW, Joo HK, Jeong KH & Lee JY (2019) Structural analysis of the manganese transport regulator MntR from Bacillus halodurans in apo and manganese bound forms. PLoS One 14, e0224689.

38 Cong X, Yuan Z, Wang Z, Wei B, Xu S & Wang J (2018) Crystal structures of manganese-dependent transcriptional repressor MntR (Rv2788) from Mycobacterium tuberculosis in apo and manganese bound forms. Biochem Biophys Res Commun 501, 423–427.

39 Posey JE, Hardham JM, Norris SJ & Gherardini FC (1999) Characterization of a manganese-dependent regulatory protein, TroR, from *Treponema pallidum*. Proceedings of the National Academy of Sciences 96, 10887–10892.

40 Olsen RJ, Sitkiewicz I, Ayeras AA, Gonulal VE, Cantu C, Beres SB, Green NM, Lei B, Humbird T, Greaver J, Chang E, Ragasa WP, Montgomery CA, Cartwright J Jr, McGeer A, Low DE, Whitney AR, Cagle PT, Blasdel TL, DeLeo FR & Musser JM (2010) Decreased necrotizing fasciitis capacity caused by a single nucleotide mutation that alters a multiple gene virulence axis. Proc Natl Acad Sci U S A 107, 888–893.

41 Toukoki C, Gold KM, McIver KS & Eichenbaum Z (2010) MtsR is a dual regulator that controls virulence genes and metabolic functions in addition to metal homeostasis in the group A streptococcus. Mol Microbiol 76, 971–989.

42 Hill PJ, Cockayne A, Landers P, Morrissey JA, Sims CM & Williams P (1998) SirR, a novel iron-dependent repressor in Staphylococcus epidermidis. Infect Immun 66, 4123–4129.

43 Rolerson E, Swick A, Newlon L, Palmer C, Pan Y, Keeshan B & Spatafora G (2006) The SloR/Dlg metalloregulator modulates Streptococcus mutans virulence gene expression. J Bacteriol 188, 5033–5044.

44 O’Rourke KP, Shaw JD, Pesesky MW, Cook BT, Roberts SM, Bond JP & Spatafora GA (2010) Genome-wide characterization of the SloR metalloregulome in Streptococcus mutans. J Bacteriol 192, 1433–1443.

45 Spatafora G, Corbett J, Cornacchione L, Daly W, Galan D, Wysota M, Tivnan P, Collins J, Nye D, Levitz T, Breyer WA & Glasfeld A (2015) Interactions of the Metalloregulatory Protein SloR from Streptococcus mutans with Its Metal Ion Effectors and DNA Binding Site. J Bacteriol 197, 3601–3615.

46 Rangachari V, Marin V, Bienkiewicz EA, Semavina M, Guerrero L, Love JF, Murphy JR & Logan TM (2005) Sequence of ligand binding and structure change in the diphtheria toxin repressor upon activation by divalent transition metals. Biochemistry 44, 5672–5682.

47 D’Aquino JA, Tetenbaum-Novatt J, White A, Berkovitch F & Ringe D (2005) Mechanism of metal ion activation of the diphtheria toxin repressor DtxR. Proc Natl Acad Sci U S A 102, 18408–18413.

48 D’Aquino JA, Lattimer JR, Denninger A, D’Aquino KE & Ringe D (2007) Role of the N-terminal helix in the metal ion-induced activation of the diphtheria toxin repressor DtxR. Biochemistry 46, 11761–11770.

49 Lavery R, Moakher M, Maddocks JH, Petkeviciute D & Zakrzewska K (2009) Conformational analysis of nucleic acids revisited: Curves+. Nucleic Acids Res 37, 5917–5929.

50 Lim S, Jung J-H, Blanchard L & de Groot A (2019) Conservation and diversity of radiation and oxidative stress resistance mechanisms in Deinococcus species. FEMS Microbiol Rev 43, 19–52.

51 Do H, Makthal N, Chandrangsu P, Olsen RJ, Helmann JD, Musser JM & Kumaraswami M (2019) Metal sensing and regulation of adaptive responses to manganese limitation by MtsR is critical for group A streptococcus virulence. Nucleic Acids Res 47, 7476–7493.

52 Jumper J, Evans R, Pritzel A, Green T, Figurnov M, Ronneberger O, Tunyasuvunakool K, Bates R, Žídek A, Potapenko A, Bridgland A, Meyer C, Kohl SAA, Ballard AJ, Cowie A, Romera-Paredes B, Nikolov S, Jain R, Adler J, Back T, Petersen S, Reiman D, Clancy E, Zielinski M, Steinegger M, Pacholska M, Berghammer T, Bodenstein S, Silver D, Vinyals O, Senior AW, Kavukcuoglu K, Kohli P & Hassabis D (2021) Highly accurate protein structure prediction with AlphaFold. Nature 596, 583–589.

53 Chakrabarti P (1990) Geometry of interaction of metal ions with histidine residues in protein structures. Protein Eng 4, 57–63.

54 Love JF, vanderSpek JC, Marin V, Guerrero L, Logan TM & Murphy JR (2004) Genetic and biophysical studies of diphtheria toxin repressor (DtxR) and the hyperactive mutant DtxR(E175K) support a multistep model of activation. Proc Natl Acad Sci U S A 101, 2506–2511.

55 Wisedchaisri G, Holmes RK & Hol WGJ (2004) Crystal structure of an IdeR-DNA complex reveals a conformational change in activated IdeR for base-specific interactions. J Mol Biol 342, 1155–1169.

56 Lee JH, Wang T, Ault K, Liu J, Schmitt MP & Holmes RK (1997) Identification and characterization of three new promoter/operators from Corynebacterium diphtheriae that are regulated by the diphtheria toxin repressor (DtxR) and iron. Infect Immun 65, 4273–4280.

57 Pohl E, Holmes RK & Hol WG (1999) Crystal structure of a cobalt-activated diphtheria toxin repressor-DNA complex reveals a metal-binding SH3-like domain. J Mol Biol 292, 653–667.

58 Pohl E, Holmes RK & Hol WG (1998) Motion of the DNA-binding domain with respect to the core of the diphtheria toxin repressor (DtxR) revealed in the crystal structures of apo- and holo-DtxR. J Biol Chem 273, 22420–22427.

59 Qiu X, Pohl E, Holmes RK & Hol WG (1996) High-resolution structure of the diphtheria toxin repressor complexed with cobalt and manganese reveals an SH3-like third domain and suggests a possible role of phosphate as co-corepressor. Biochemistry 35, 12292–12302.

60 Stevenson B, Wyckoff EE & Payne SM (2016) Vibrio cholerae FeoA, FeoB, and FeoC Interact To Form a Complex. J Bacteriol 198, 1160–1170.

61 Marcos-Torres FJ, Juniar L & Griese JJ (2023) The molecular mechanisms of the bacterial iron sensor IdeR. Biochem Soc Trans 51, 1319–1329.

62 de Sanctis D, Beteva A, Caserotto H, Dobias F, Gabadinho J, Giraud T, Gobbo A, Guijarro M, Lentini M, Lavault B, Mairs T, McSweeney S, Petitdemange S, Rey-Bakaikoa V, Surr J, Theveneau P, Leonard GA & Mueller-Dieckmann C (2012) ID29: a high-intensity highly automated ESRF beamline for macromolecular crystallography experiments exploiting anomalous scattering. J Synchrotron Radiat 19, 455–461.

63 Nurizzo D, Mairs T, Guijarro M, Rey V, Meyer J, Fajardo P, Chavanne J, Biasci JC, McSweeney S & Mitchell E (2006) The ID23-1 structural biology beamline at the ESRF. J Synchrotron Radiat 13, 227–238.

64 Kabsch W (2010) XDS. Acta Crystallogr D Biol Crystallogr 66, 125–132.

65 Sheldrick GM (2008) A short history of \it SHELX. Acta Crystallogr A 64, 112–122.

66 Morris RJ, Perrakis A & Lamzin VS (2003) ARP/wARP and automatic interpretation of protein electron density maps. Methods Enzymol 374, 229–244.

67 Emsley P & Cowtan K (2004) Coot: model-building tools for molecular graphics. Acta Crystallogr D Biol Crystallogr 60, 2126–2132.

68 Adams PD, Afonine PV, Bunkóczi G, Chen VB, Davis IW, Echols N, Headd JJ, Hung L-W, Kapral GJ, Grosse-Kunstleve RW, McCoy AJ, Moriarty NW, Oeffner R, Read RJ, Richardson DC, Richardson JS, Terwilliger TC & Zwart PH (2010) PHENIX: a comprehensive Python-based system for macromolecular structure solution. Acta Crystallogr D Biol Crystallogr 66, 213–221.

69 McCoy AJ, Grosse-Kunstleve RW, Adams PD, Winn MD, Storoni LC & Read RJ (2007) Phaser crystallographic software. J Appl Crystallogr 40, 658–674.

70 Pernot P, Round A, Barrett R, De Maria Antolinos A, Gobbo A, Gordon E, Huet J, Kieffer J, Lentini M, Mattenet M, Morawe C, Mueller-Dieckmann C, Ohlsson S, Schmid W, Surr J, Theveneau P, Zerrad L & McSweeney S (2013) Upgraded ESRF BM29 beamline for SAXS on macromolecules in solution. J Synchrotron Radiat 20, 660–664.

71 Franke D, Petoukhov MV, Konarev PV, Panjkovich A, Tuukkanen A, Mertens HDT, Kikhney AG, Hajizadeh NR, Franklin JM, Jeffries CM & Svergun DI (2017) ATSAS 2.8: a comprehensive data analysis suite for small-angle scattering from macromolecular solutions. J Appl Crystallogr 50, 1212–1225.

72 Petoukhov MV, Franke D, Shkumatov AV, Tria G, Kikhney AG, Gajda M, Gorba C, Mertens HDT, Konarev PV & Svergun DI (2012) New developments in the ATSAS program package for small-angle scattering data analysis. J Appl Crystallogr 45, 342–350.

73 Manalastas-Cantos K, Konarev PV, Hajizadeh NR, Kikhney AG, Petoukhov MV, Molodenskiy DS, Panjkovich A, Mertens HDT, Gruzinov A, Borges C, Jeffries CM, Svergun DI & Franke D (2021) ATSAS 3.0: expanded functionality and new tools for small-angle scattering data analysis. J Appl Crystallogr 54, 343–355.

74 Guinier A & Fournet G (1955) Small-angle Scattering of X-rays Wiley.

75 Svergun DI (1992) Determination of the regularization parameter in indirect-transform methods using perceptual criteria. J Appl Crystallogr 25, 495–503.

76 Franke D & Svergun DI (2009) DAMMIF, a program for rapid ab-initio shape determination in small-angle scattering. J Appl Crystallogr 42, 342–346.

77 Volkov VV, Svergun DI & IUCr (2003) Uniqueness of ab initio shape determination in small-angle scattering. J Appl Crystallogr 36, 860–864.

78 Luscombe NM, Laskowski RA & Thornton JM (1997) NUCPLOT: A Program to Generate Schematic Diagrams of Protein-Nucleic Acid Interactions. Nucleic Acids Res 25, 4940–4945.

