## Supplementary Material for "Metal ion activation and DNA recognition by the *Deinococcus radiodurans* manganese sensor DR2539"

#### Tables

Table S1 - Crystallographic data collection and refinement statistics

| **Structure** | **Cd-DR2539 (SAD)** | **Cd-DR2539 (MR)** | **Mn-DR2539-*dr1709p*** |
|---|---|---|---|
| **PDB ID** | 8PVT | 8PVZ | 8PW0 |
| **Data collection and processing** | | | |
| Beamline | ID29 (ESRF) | ID23-1 (ESRF) | ID29 (ESRF) |
| Wavelength | 1.7712 | 0.9793 | 0.9762 |
| Resolution (Å) | 43.60 - 2.50 (2.64 - 2.50) | 46.92 - 2.00 (2.11 - 2.00) | 47.88 - 2.20 (2.32 - 2.20) |
| Space group | $P4_32_12$ | $P4_32_12$ | $P4_22_12$ |
| Cell dimensions a, b, c (Å) | 60.846, 60.846, 187.49 | 60.785, 60.785, 187.681 | 67.710, 67.710, 158.137 |
| No. unique reflections | 12916 (1752) | 24727 (3534) | 19507 (2768) |
| Rmerge | 0.168 (1.014) | 0.103 (1.221) | 0.093 (1.016) |
| Rmeas | 0.187 (1.074) | 0.113 (1.344) | 0.099 (1.074) |
| Rpim | 0.038 (0.256) | 0.047 (0.555) | 0.033 (0.339) |
| I/sigmaI | 16.2 (2.8) | 11.4 (2.2) | 13.1 (1.9) |
| Completeness (%) | 99.4 (96.3) | 99.8 (99.9) | 100.0 (100.0) |
| Multiplicity | 23.2 (16.5) | 5.5 (5.8) | 8.8 (9.2) |
| $CC_{1/2}$ | 0.998 (0.864) | 0.998 (0.622) | 0.998 (0.678) |
| Anomalous completeness (%) | 99.6 (96.3) | 97.3 (99.2) | 99.98 (99.8) |
| Anomalous multiplicity | 23.2 (16.5) | 2.9 (2.9) | 4.7 (4.8) |
| **Refinement** | | | |

| | | | |
|---|---|---|---|
| Resolution (Å) | 43.59 - 2.5 (2.60 - 2.50) | 43.60 - 2.00 (2.08 - 2.00) | 47.88 - 2.20 (2.26 - 2.20) |
| Nº of reflections | 12852 | 24643 | 19457 |
| Rwork/Rfree (%)[a] | 20.49/25.50 | 18.56/22.00 | 18.26/22.73 |
| **Nº of non H- atoms** | | | |
| Protein | 2078 | 2093 | 1046 |
| DNA | - | - | 856 |
| Metal ions | 23 | 24 | 4 |
| Water | 57 | 139 | 92 |
| **B-factors** | | | |
| Protein (Å$^2$) | 40.00 | 41.00 | 58.23 |
| DNA (Å$^2$) | - | | 50.96 |
| Metal ions (Å$^2$) | 77.66 | 55.78 | 81.11 |
| **R.m.s. deviations** | | | |
| Bond lengths (Å) | 0.004 | 0.013 | 0.008 |
| Bond angles (Å) | 0.706 | 1.185 | 1.095 |
| Ramachadran favored/poor (%)[b] | 99.25/0.75 | 99.26/0.74 | 97.78/1.48 |
| Rama *Z*-score[b] | -1.01 | 0.91 | -0.38 |
| Clashscore[b] | 6.96 | 3.57 | 3.20 |
| Molprobity score[b] | 1.67 | 1.28 | 1.36 |

Values in parentheses are for the highest resolution shell. Friedel pairs were merged.

[a]$R_{free}$ is calculated from a randomly selected subset of ~5% of reflections excluded from refinement.

[b]Geometry statistics were calculated with MolProbity [1].

Table S2 - Dimer interface of DR2539 truncated form.

| Hydrogen bonds | | | Salt bridges | | |
|---|---|---|---|---|---|
| Monomer A | Distance (Å) | Monomer B | Monomer A | Distance (Å) | Monomer B |
| B:SER 109[ OG ] | 2.82 | A:GLU 100[ OE1] | B:ARG 111[ NE ] | 3.36 | A:GLU 96[ OE1] |
| B:ARG 111[ NE ] | 3.18 | A:GLU 100[ OE2] | B:ARG 111[ NE ] | 3.18 | A:GLU 100[ OE2] |
| B:ARG 111[ NH2] | 3.05 | A:GLU 100[ OE2] | B:ARG 111[ NH1] | 3.66 | A:GLU 96[ OE1] |
| B:ARG 111[ NH2] | 3.81 | A:ASP 99[ OD2] | B:ARG 111[ NH2] | 3.05 | A:GLU 100[ OE2] |
| B:ARG 115[ NE ] | 3.66 | A:GLU 96[ OE1] | B:ARG 111[ NH2] | 3.81 | A:ASP 99[ OD2] |
| B:ARG 115[ NH1] | 3.90 | A:GLY 91[ O ] | B:ARG 115[ NE ] | 3.66 | A:GLU 96[ OE1] |
| B:ARG 115[ NH2] | 3.77 | A:GLU 96[ OE2] | B:ARG 115[ NH2] | 3.77 | A:GLU 96[ OE2] |
| B:ARG 115[ NH2] | 3.70 | A:GLU 96[ OE1] | B:ARG 115[ NH2] | 3.70 | A:GLU 96[ OE1] |
| B:TRP 119[ NE1] | 3.23 | A:ALA 89[ O ] | B:GLU 96[ OE1] | 3.18 | A:ARG 115[ NH2] |
| B:ALA 89[ O ] | 3.69 | A:TRP 119[ NE1] | B:GLU 96[ OE1] | 3.51 | A:ARG 115[ NE ] |
| B:GLU 96[ OE1] | 3.18 | A:ARG 115[ NH2] | B:GLU 96[ OE2] | 3.52 | A:ARG 115[ NH2] |
| B:GLU 96[ OE1] | 3.51 | A:ARG 115[ NE ] | - | - | - |
| B:GLU 100[ OE2] | 2.67 | A:SER 109[ OG ] | - | - | - |

**Table S3-** Cadmium atoms in structure 8PVZ.

| Ion | Occ | B-factor (Å$^2$) |
|---|---|---|
| CD1 | 1.00 | 34.12 |
| CD2 | 1.00 | 34.63 |
| CD3 | 1.00 | 33.13 |
| CD4 | 1.00 | 35.22 |
| CD5 | 0.82 | 45.61 |
| CD6 | 0.80 | 42.68 |
| CD7 | 0.89 | 36.08 |
| CD8 | 0.83 | 33.29 |
| CD9 | 0.92 | 34.89 |
| CD10 | 0.91 | 58.04 |
| CD11 | 0.93 | 48.04 |
| CD12 | 0.88 | 49.74 |

| | | |
|---|---|---|
| CD13 | 0.30 | 47.44 |
| CD14 | 0.46 | 76.98 |
| CD15 | 0.57 | 92.76 |
| CD16 | 0.60 | 49.38 |
| CD17 | 0.96 | 47.56 |
| CD18 | 0.59 | 126.55 |
| CD19 | 0.58 | 93.27 |
| CD20 | 0.29 | 60.08 |
| CD21A | 0.51 | 55.73 |
| CD21B | 0.49 | 64.46 |
| CD22A | 0.54 | 81.27 |
| CD22B | 0.46 | 57.91 |

**Table S4-** Protein-DNA interface.

| DNA | Distance (Å) | Monomer A |
|---|---|---|
| B:DG 12[ OP1] | 2.46 | A:SER 6[ OG ] |
| B:DC 13[ OP1] | 2.87 | A:SER 8[ OG ] |
| B:DC 13[ OP2] | 3.15 | A:TYR 12[ OH ] |
| B:DT 5[ OP2] | 2.48 | A:THR 27[ OG1] |
| B:DT 4[ OP2] | 2.80 | A:GLN 28[ N ] |
| B:DC 14[ OP2] | 2.91 | A:ALA 37[ N ] |
| B:DC 14[ OP2] | 2.57 | A:SER 40[ OG ] |
| B:DT 5[ OP2] | 2.81 | A:THR 42[ OG1] |
| B:DT 6[ OP2] | 3.34 | A:ARG 46[ NH1] |
| B:DG 12[ OP2] | 2.52 | A:LYS 47[ NZ ] |
| B:DT 5[ OP1] | 2.80 | A:HIS 56[ NE2] |
| B:DT 5[ OP1] | 2.89 | A:TYR 59[ N ] |
| B:DT 4[ O2 ] | 3.62 | A:TYR 59[ OH ] |

Table S5- Oligonucleotides sequences of the *dr1709* promotor region and variants used for crystallization and EMSA assays. For sake of readability, forward primers are shown 5’ to 3’ and reverse 3’ to 5’.

| Oligo name | Primers | Sequences |
|---|---|---|
| dr1709p_wt | dr1709p_wt_for<br>dr1709p_wt_rev | ATTTTAGTCGCGCCTAAAATA<br>TAAAATCAGCGCGGATTTTAT |
| dr1709p_T2G | dr1709p_T2G_for<br>dr1709p_T2G_rev | A**G**TTTAGTCGCGCCTAAA**C**TA<br>T**C**AAATCAGCGCGGATTT**G**AT |
| dr1709p_T3G | dr1709p_T3G_for<br>dr1709p_T3G_rev | AT**G**TTAGTCGCGCCTAA**C**ATA<br>TA**C**AATCAGCGCGGATT**G**TAT |
| dr1709p_T4G | dr1709p_T4G_for<br>dr1709p_T4G_rev | ATT**G**TAGTCGCGCCTA**C**AATA<br>TAA**C**ATCAGCGCGGAT**G**TTAT |
| dr1709p_T5G | dr1709p_T5G_for<br>dr1709p_T5G_rev | ATTT**G**AGTCGCGCCT**C**AAATA<br>TAAA**C**TCAGCGCGGA**G**TTTAT |
| dr1709p_T2345G | dr1709p_T2345G_for<br>dr1709p_T2345G_rev | A**GGGG**AGTCGCGCCT**CCCC**TA<br>T**CCCC**TCAGCGCGGA**GGGG**AT |
| dr1709p_T345G | dr1709p_T345G_for<br>dr1709p_T345G_rev | AT**GGG**AGTCGCGCCT**CCC**ATA<br>TA**CCC**TCAGCGCGGA**GGG**TAT |
| dr1709p_T2C | dr1709p_T2C_for<br>dr1709p_T2C_rev | A**C**TTTAGTCGCGCCTAAA**G**TA<br>T**G**AAATCAGCGCGGATTT**C**AT |
| dr1709p_T4C | dr1709p_T4C_for<br>dr1709p_T4C_rev | ATT**C**TAGTCGCGCCTA**G**AATA<br>TAA**G**ATCAGCGCGGAT**C**TTAT |
| dr1709p_A6G | dr1709p_A6G_for<br>dr1709p_A6G_rev | ATTTT**G**GTCGCGCC**C**AAAATA<br>TAAAA**C**CAGCGCGG**G**TTTTAT |
| dr1709p_T4C/A6G | dr1709p_T4C/A6G_for<br>dr1709p_T4C/A6G_rev | ATT**C**T**G**GTCGCGCC**C**A**G**AATA<br>TAA**G**A**C**CAGCGCGG**G**T**C**TTAT |

Table S6 - Small angle X-ray scattering data collection and processing.

| **Data-collection parameters** | | | | | | |
|---|---|---|---|---|---|---|
| **DR2539 form** | Full length | Full length | Full length | Truncate d | Truncate d | Truncate d |

| Condition | as-isolated | Mn | EDTA | DR2539tr - as-isolated | DR2539tr - Mn | DR2539tr - EDTA |
|---|---|---|---|---|---|---|
| **Instrument** | ESRF BM29 | ESRF BM29 | ESRF BM29 | ESRF BM29 | ESRF BM29 | ESRF BM29 |
| **Wavelength (Å)** | 0.9919 | 0.9919 | 0.9919 | 0.9919 | 0.9919 | 0.9919 |
| **q-range (Å-1)** | 0.0033-0.49 | 0.0033-0.49 | 0.0035-0.50 | 0.0035-0.50 | 0.0035-0.50 | 0.0035-0.50 |
| **Sample-to-detector distance (m)** | 2.872 | 2.872 | 2.867 | 2.867 | 2.867 | 2.867 |
| **Exposure time (sec)** | 10 ’ 4 | 10 ’ 4 | 15 ’ 1 | 10 ’ 1 | 10 ’ 1 | 10 ’ 1 |
| **Concentration range (mg/mL)** | 1.72 – 3.44 | 0.97 – 3.87 | 1.13 – 6.50 | 0.25 – 5 | 0.25 – 9.04 | 0.25 – 10.38 |
| **Temperature (K)** | 293 | 293 | 293 | 293 | 293 | 293 |
| **Detector** | Pilatus 1M (Dectris) | Pilatus 1M (Dectris) | Pilatus 1M (Dectris) | Pilatus 1M (Dectris) | Pilatus 1M (Dectris) | Pilatus 1M (Dectris) |
| **Beam size (µm2)** | 700 x 700 | 700 x 700 | 700 x 700 | 700 x 700 | 700 x 700 | 700 x 700 |
| **Data processing** | | | | | | |
| **I0 (cm-1) [from Guinier]** | 0.03103 | 0.03075 | 0.02915 | 0,02517 | 0,02988 | 0,02576 |
| **Rg (nm) [from Guinier]** | 2.84 | 2.85 | 2.86 | 2.67 | 2.46 | 2.78 |
| **qminRg – qmaxRg used for Guinier** | 0.44 -1.30 | 0.35 – 1.29 | 0.42 – 1.27 | 0.27 – 1.30 | 0.38 – 1.29 | 0.36 - 1.29 |
| **Dmax (Å) [from p(r) ]** | 82.4 | 86.2 | 102.9 | 86.7 | 69.1 | 87.0 |
| **q-range used for p(r) (Å-1)** | 0.015 – 0.28 | 0.012 – 0.28 | 0.015 – 0.28 | 0.010 - 0.30 | 0.015 - 0.32 | 0.013 – 0.29 |
| **Porod volume Vp (Å3) [from Scatter]** | 81602 | 68479 | 79011 | 49355 | 45384 | 54074 |

| | | | | | | |
|---|---|---|---|---|---|---|
| **Molecular mass Mr (kDa) [from Vp]** | 48.001 | 40.281 | 46.477 | 29.032 | 26.696 | 31.808 |
| **Calculated dimeric Mr from sequence (kDa)** | 49.088 | 49.088 | 49.088 | 31796 | 31796 | 31796 |

### Figures

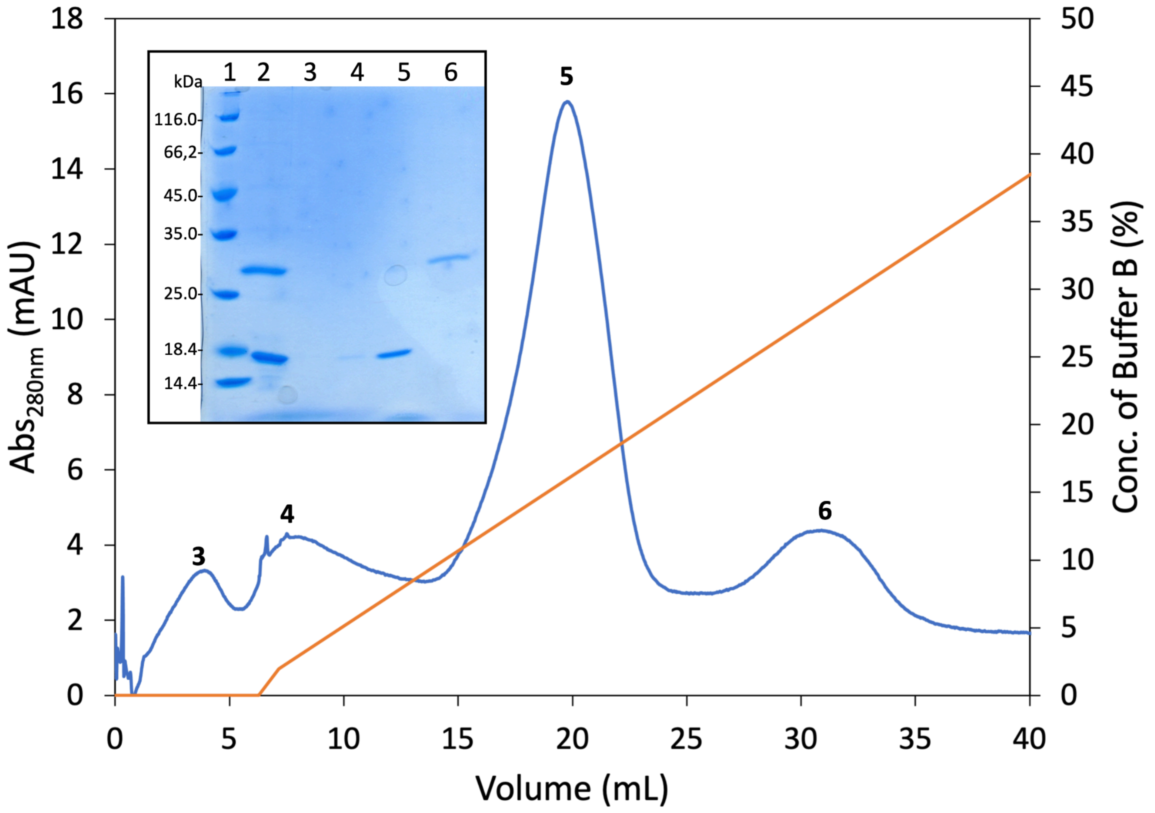


**Figure S1 -** HiTrap Heparin HP affinity chromatography profile separation of truncated and full length forms of DR2539. Absorbance at 280 nm is shown in blue and the concentration of buffer B is represented in orange. Inset: SDS-PAGE of fractions collected during purification 1) Molecular weight marker (Thermo Fisher Scientific); 2) DR2539 sample before injection; 3) Peak 3 of the chromatogram; 4) Peak 4 of the chromatogram; 3) Peak 5 of the chromatogram; 4) Peak 6 of the chromatogram.

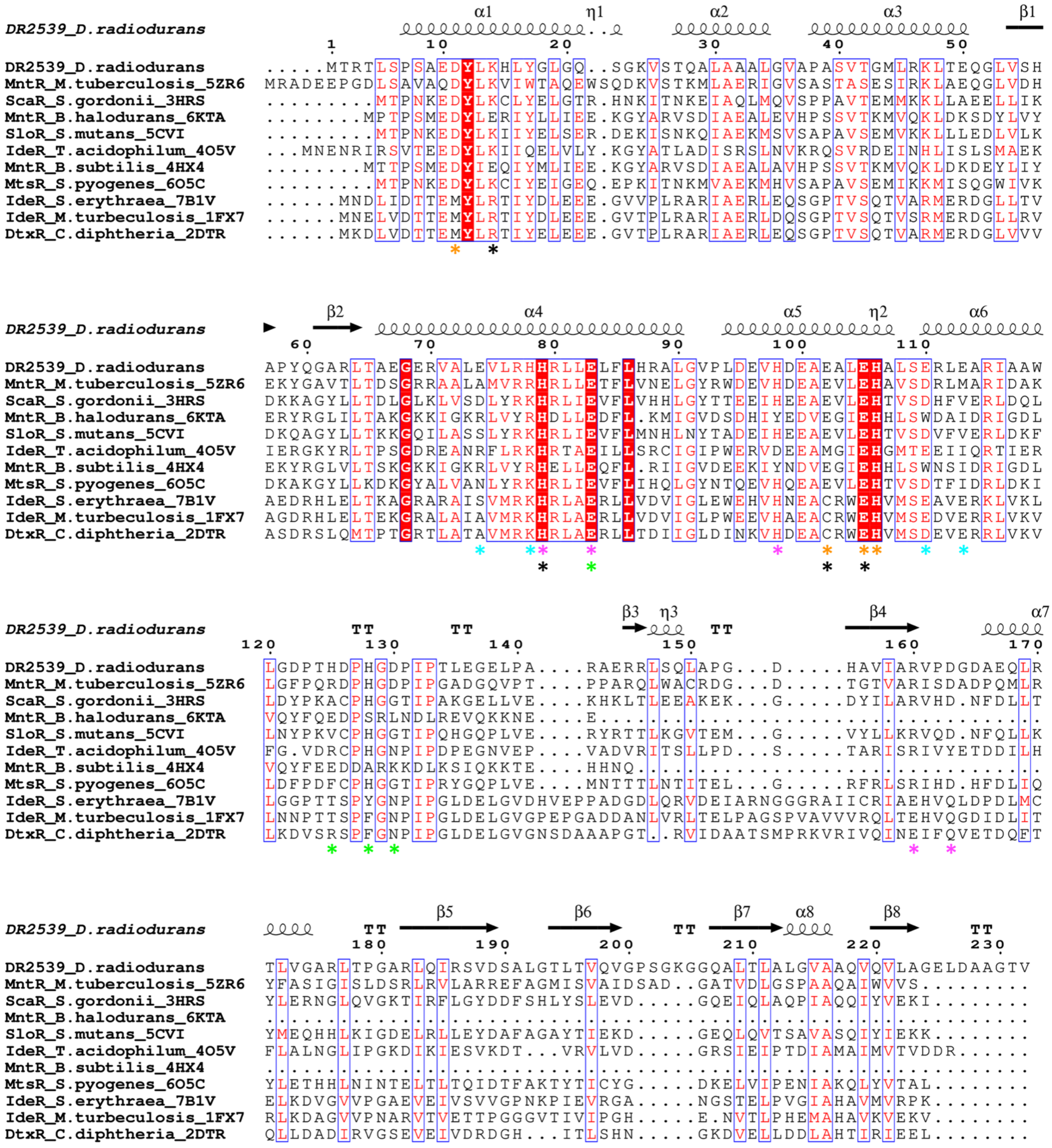


**Figure S2** - Sequence alignment of *D. radiodurans* DR2539 and representatives of DtxR/MntR family. Metal binding amino acids of all members marked with asterisks: primary site in orange, secondary site in black (present in MntRs), ancillary site I and II in magenta and green, respectively, and tertiary site in cyan. Representation generated by ESPript 3.0

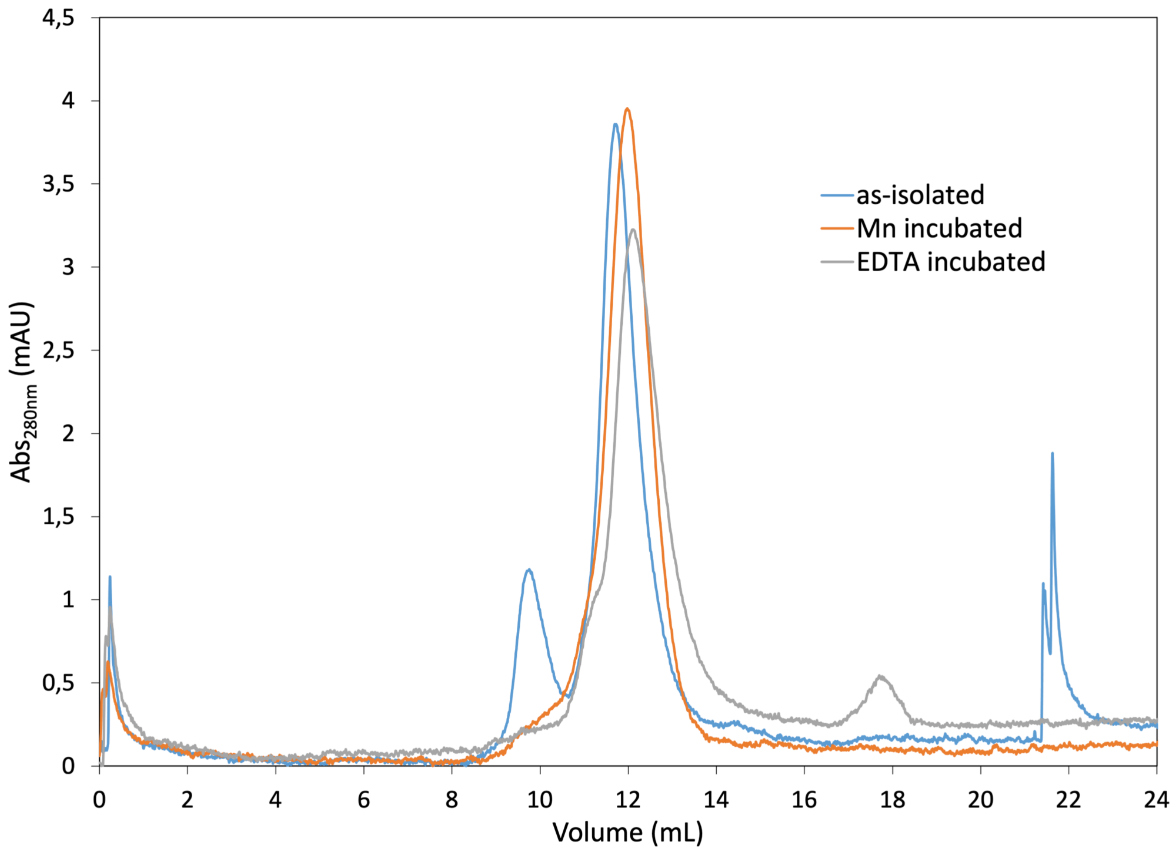


**Figure S3** – Superdex S75 elution profile of DR2539 as-isolated (blue line) and incubated with EDTA (grey line) and manganese (orange line).

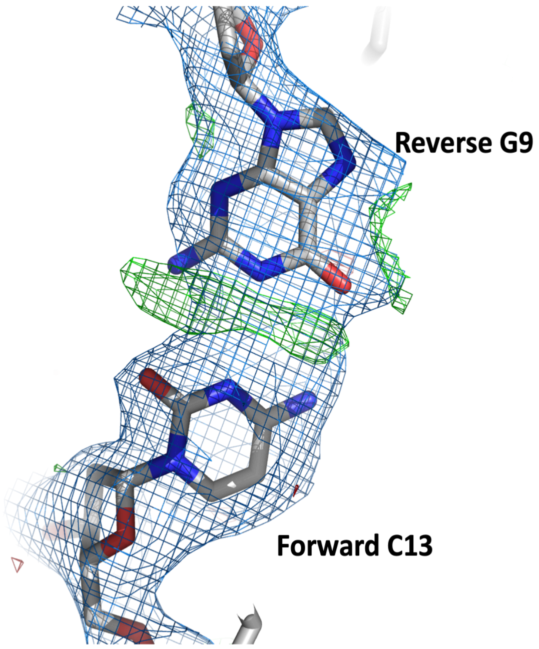


**Figure S4** - DNA maps ambiguity. Due to the quasi-palindromic symmetry and the 2-fold crystallographic axis these nucleotide pairs are modelled in double conformation to reflect the two possible sequences. 2Fo-Fc (blue mesh) and Fo-Fc (green) electron density maps contoured at 2σ and 3σ, respectively.

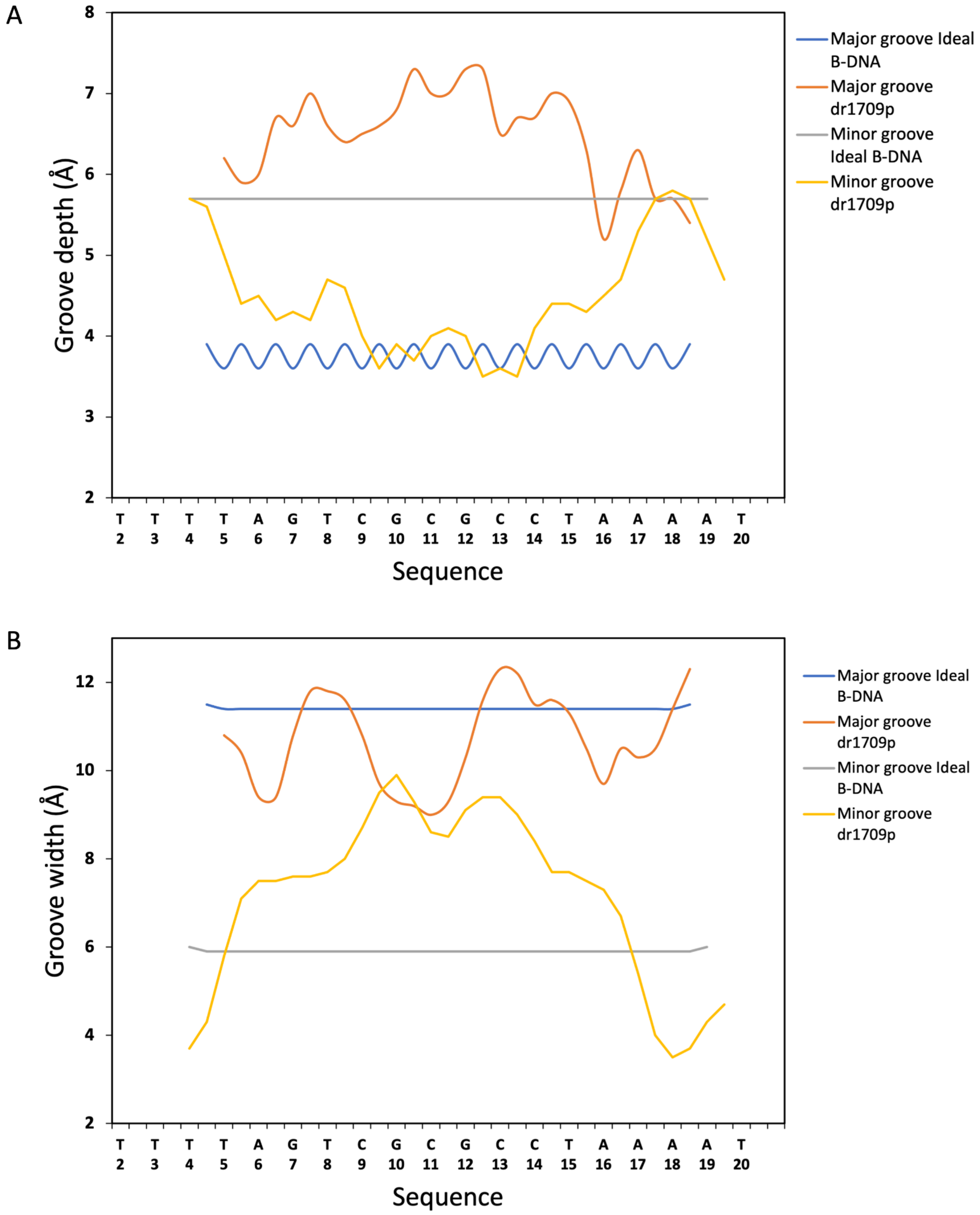


**Figure S5** – Major and minor grooves parameters calculated by Curves+ for *dr1709p* and ideal B-DNA [2].

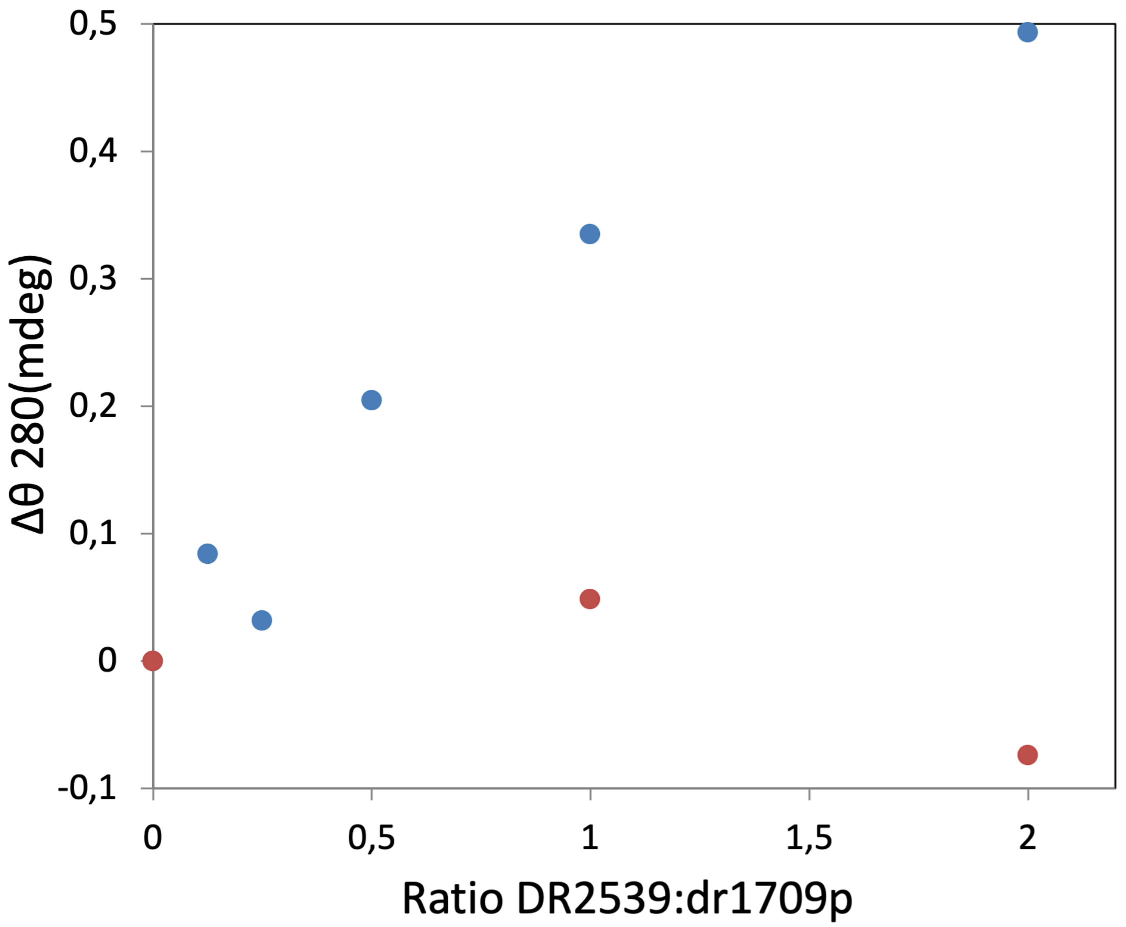


**Figure S6** – CD spectroscopy analysis. Binding of DR2539 to *dr1709p* was monitored by circular dichroism (CD) spectroscopy. DR2539 protein was added incrementally to *dr1709p* (2 µM) in the binding buffer (blue dots) and a metal depletion test was performed in the buffer supplemented with 2 mM EDTA (orange dots).

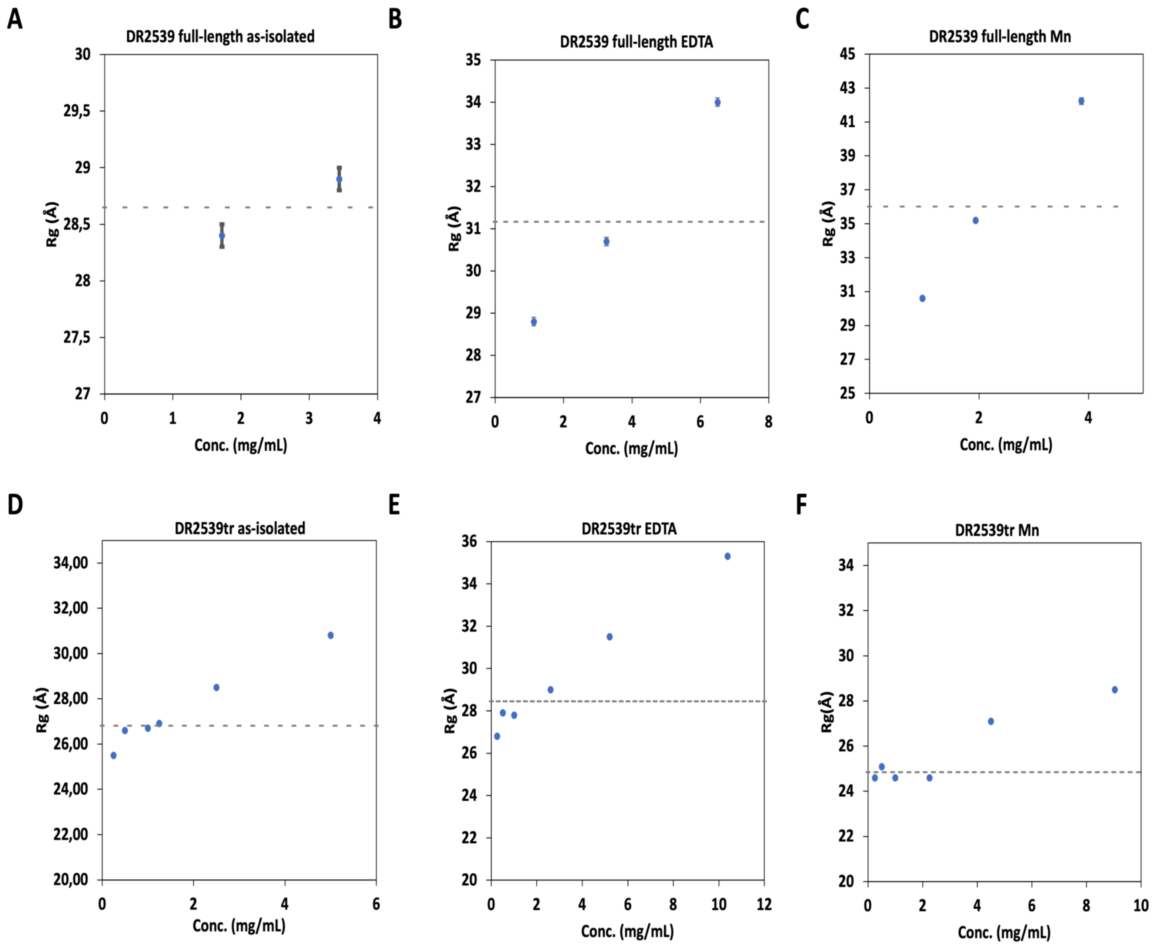


**Figure S7** - Small angle X-ray scattering (SAXS) of DR2539 full-length and truncated. The Guinier $R_g$ values are calculated to the several protein concentrations in three different conditions (as isolated, EDTA incubated and Mn incubated). Average Rg value is represented by the dashed line.

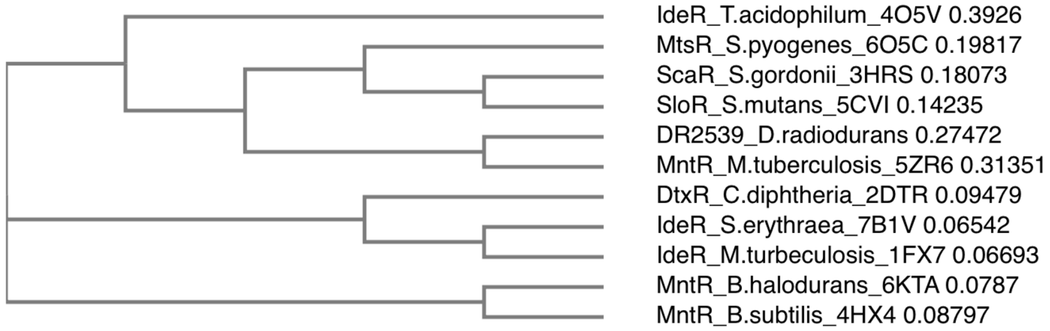


**Figure S8** - Phylogenetic tree. Comparison of DBDs sequences from the DtxR family members (Clustal Omega [3]).

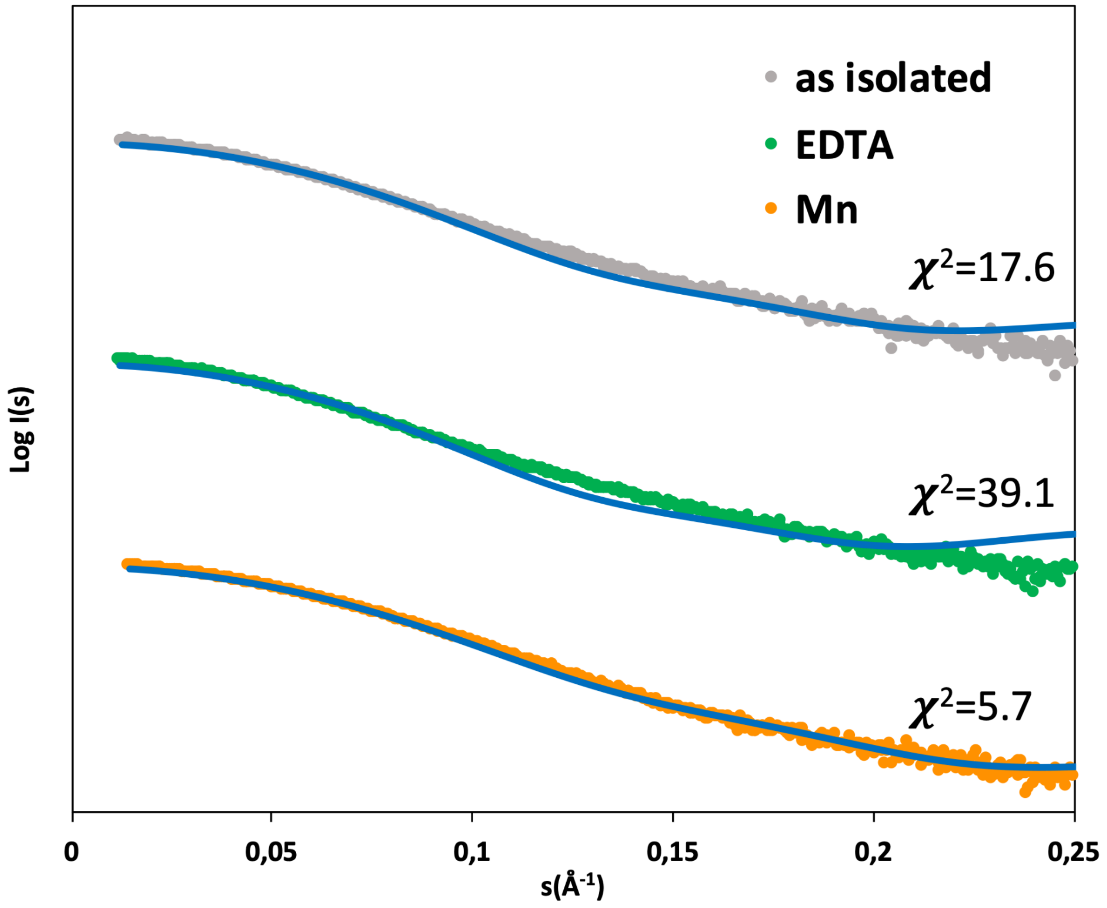


**Figure S9 -** Comparison of the experimental scattering curves of truncated DR2539 as isolated (grey dots) and incubated with EDTA (green dots) or manganese (orange dots). Theoretical scattering curves (blue lines) from the crystal structure (PDB ID 8PW0) calculated by CRYSOL.

**References**


1 Davis IW, Leaver-Fay A, Chen VB, Block JN, Kapral GJ, Wang X, Murray LW, Arendall WB 3rd, Snoeyink J, Richardson JS & Richardson DC (2007) MolProbity: all-atom contacts and structure validation for proteins and nucleic acids. *Nucleic Acids Res* **35**, W375–83.

2 Lavery R, Moakher M, Maddocks JH, Petkeviciute D & Zakrzewska K (2009) Conformational analysis of nucleic acids revisited: Curves+. *Nucleic Acids Res* **37**, 5917–5929.

3 Sievers F, Wilm A, Dineen D, Gibson TJ, Karplus K, Li W, Lopez R, McWilliam H, Remmert M, Söding J, Thompson JD & Higgins DG (2011) Fast, scalable generation of high-quality protein multiple sequence alignments using Clustal Omega. *Mol Syst Biol* **7**, 539.
